# Regression and Alignment for Functional Data and Network Topology

**DOI:** 10.1101/2023.07.13.548836

**Authors:** Danni Tu, Julia Wrobel, Theodore D. Satterthwaite, Jeff Goldsmith, Ruben C. Gur, Raquel E. Gur, Jan Gertheiss, Dani S. Bassett, Russell T. Shinohara

**Affiliations:** The Penn Statistics in Imaging and Visualization Endeavor (PennSIVE), Department of Biostatistics, Epidemiology, and Informatics, University of Pennsylvania, Philadelphia, PA, USA; Department of Biostatistics and Bioinformatics, Emory University, Atlanta, GA, USA; Department of Psychiatry, Perelman School of Medicine, Philadelphia, PA USA; Penn Lifespan Informatics and Neuroimaging Center, Philadelphia, PA, USA; Department of Biostatistics, Columbia University, New York, NY, USA; The Penn Medicine-CHOP Lifespan Brain Institute, Philadelphia, PA, USA; Department of Mathematics and Statistics, School of Economics and Social Sciences, Helmut Schmidt University, Hamburg, Germany; Department of Bioengineering, University of Pennsylvania, Philadelphia, PA, USA; Department of Physics and Astronomy, University of Pennsylvania, Philadelphia, PA, USA; Department of Electrical and Systems Engineering, University of Pennsylvania, Philadelphia, PA, USA; Department of Neurology, University of Pennsylvania, Philadelphia, PA, USA

**Keywords:** Alignment, Functional Data Analysis, Functional Regression, Network Neuroscience

## Abstract

In the brain, functional connections form a network whose topological organization can be described by graph-theoretic network diagnostics. These include characterizations of the community structure, such as modularity and participation coefficient, which have been shown to change over the course of childhood and adolescence. To investigate if such changes in the functional network are associated with changes in cognitive performance during development, network studies often rely on an arbitrary choice of pre-processing parameters, in particular the proportional threshold of network edges. Because the choice of parameter can impact the value of the network diagnostic, and therefore downstream conclusions, we propose to circumvent that choice by conceptualizing the network diagnostic as a function of the parameter. As opposed to a single value, a network diagnostic curve describes the connectome topology at multiple scales—from the sparsest group of the strongest edges to the entire edge set. To relate these curves to executive function and other covariates, we use scalar-on-function regression, which is more flexible than previous functional data-based models used in network neuroscience. We then consider how systematic differences between networks can manifest in misalignment of diagnostic curves, and consequently propose a supervised curve alignment method that incorporates auxiliary information from other variables. Our algorithm performs both functional regression and alignment via an iterative, penalized, and nonlinear likelihood optimization. The illustrated method has the potential to improve the interpretability and generalizability of neuroscience studies where the goal is to study heterogeneity among a mixture of function- and scalar-valued measures.

## 1. Introduction

Functional magnetic resonance imaging (fMRI) is a powerful, non-invasive method to measure blood oxygen level dependent (BOLD) signals in the brain. A major goal of network neuroscience studies is to use the observed coherence of these BOLD time series to uncover functional relationships between brain regions. These connections, which collectively form a functional network or *connectome*, exhibit a complex hierarchical and small-world topology that promotes efficient information processing (Muldoon *and others*, 2016) and can be described by various graph-theoretical diagnostics (Bullmore and Bassett, 2011).

Importantly, the functional connectome is modular: it can be partitioned into communities that are strongly intraconnected and relatively weakly interconnected (Garcia *and others*, 2018; Sporns and Betzel, 2016). These communities are either known *a priori* or may be estimated from the data using greedy algorithms (Fortunato, 2010; Gates *and others*, 2016). The modularity index quantifies the saturation of within-community connections relative to that of a null graph where community structure has been reconfigured at random (Newman and Girvan, 2004). Relatedly, the participation coefficient (Betzel and Bassett, 2017) assesses the degree to which nodes are connected to different communities. Modular systems and other characterizations of mesoscale network organization have been linked to cognitive outcomes in both healthy adults (Gallen and D’Esposito, 2019) and neurodegenerative disease (Gamboa *and others*, 2014). In adolescents, reorganization and integration of modules during development (Fair *and others*, 2009; Gu *and others*, 2019; Morgan *and others*, 2018) are thought to support simultaneous changes in executive functioning (Baum *and others*, 2017).

In most connectome studies, the matrix corresponding to network edge weights is thresholded (i.e., sparsified) to limit the data to connections with stronger signal (Wang *and others*, 2009; Geerligs *and others*, 2014; Achard, 2006). However, there is much disagreement about what threshold to use, and this somewhat arbitrary choice can significantly impact the conclusions about how network structure relates to group differences and other outcomes (Garrison *and others*, 2015; Scheinost *and others*, 2012). We circumvent this choice by considering a range of all possible thresholds *t* (Bullmore and Bassett, 2011; Bassett *and others*, 2012). As a result, the network diagnostics—modularity and participation coefficient—are functions of *t* (Figure 1). To relate these functional data to scalar-valued outcomes (e.g., executive function) and covariates (e.g., age and motion), we introduce a functional regression framework.

**Fig. 1.**
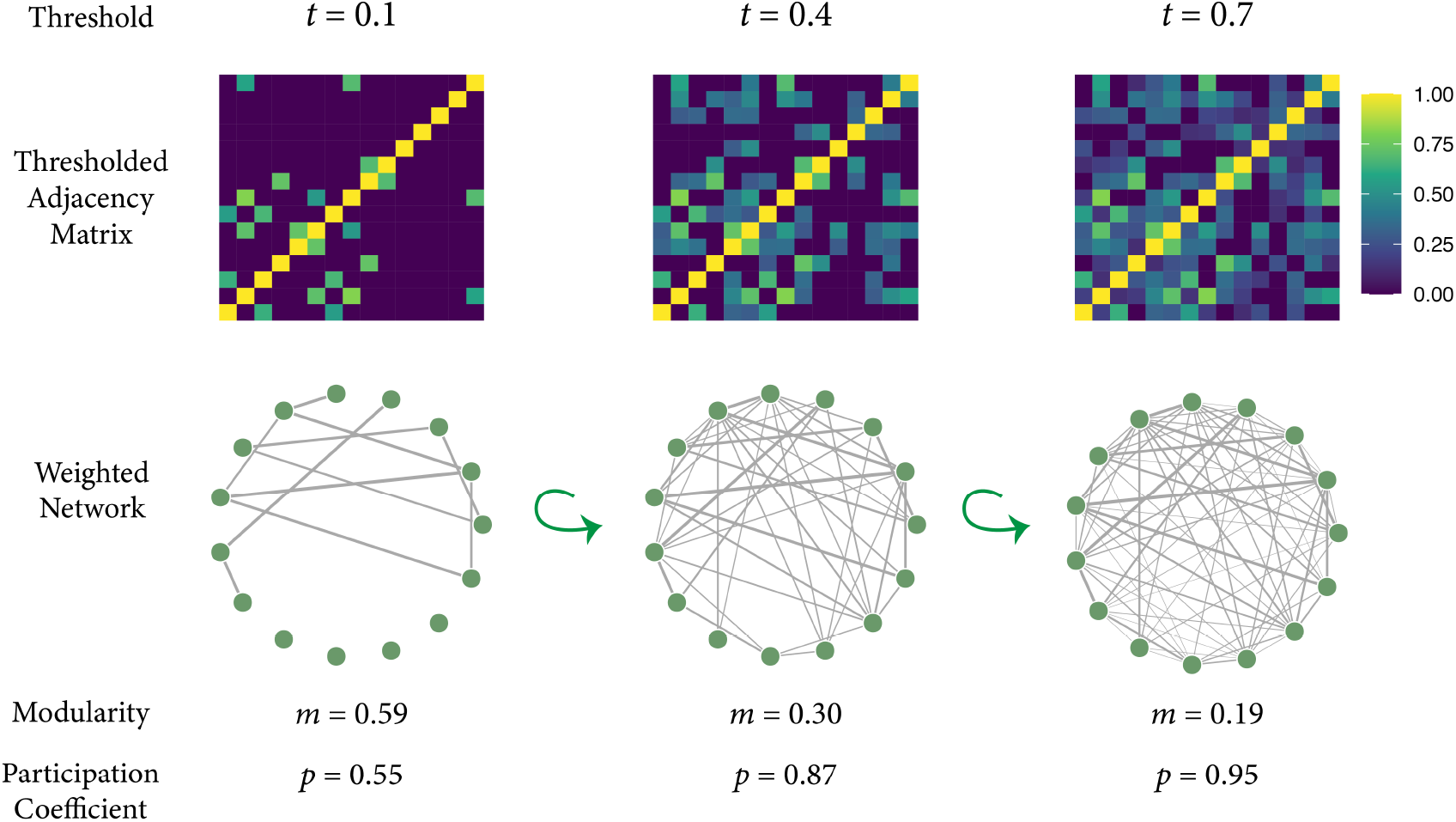
Network diagnostic curves represent graph topology at multiple scales. For any adjacency matrix, the threshold *t* ∈ [0, 1] represents the proportion of strongest edges to keep, which defines a sparser weighted network. When *t* = 0, the resulting network is the set of disconnected nodes. As *t* increases, edges are added to those nodes in decreasing order of strength. For each network, various network diagnostics describing the topology of edge connections can be evaluated (e.g., modularity and participation coefficient). Therefore, calculating the network diagnostic at multiple values of *t* ∈ [0, 1] results in a network diagnostic curve.

Our proposed statistical model, Regression and Alignment for Functional Data and Network Topology (RAFT), is motivated by three aspects of the data that current methods for network diagnostic curves do not address. First, we address the function-valued nature of our data by using a scalar-on-function regression model, which is more powerful than pointwise comparisons of curves at fixed values of *t* (Figure S.7). Regression models can be used to predict a continuous scalar-valued outcome, in contrast to permutation tests which are more often used in neuroscience studies (Bassett *and others*, 2012; Singh *and others*, 2013; Palaniyappan *and others*, 2016). And through functional principal components analysis (FPCA), we can describe variation in the curve shape that cannot be evaluated by existing summaries such as the area under the curve (Giusti *and others*, 2015). Second, our method addresses misalignment in the functional data, which can materialize due to confounding in network diagnostic measures that stems from systematic differences in edge strength distribution (van den Heuvel *and others*, 2017). To align the data, we introduce a simple parameterization of a warping function based on the cumulative distribution of a Beta random variable. Third, while the majority of alignment methods use only the information observed in the curves themselves (“unsupervised” alignment), we propose a *supervised* algorithm that incorporates information from other covariates.

In simulated data, we demonstrate that our proposed curve alignment methods improve parameter estimation in downstream regression analyses. We then apply our method to network and neurocognitive data obtained from the Philadelphia Neurodevelopmental Cohort (PNC), a community-based study of brain development in youth aged 8 to 21 years (Satterthwaite *and others*, 2016), thereby complementing previous findings that link the differentiation of cortical systems to cognitive performance in the PNC (Gu *and others*, 2015).

## 2. Functional Data Analysis of Multiscale Network Topology

### 2.1. Brain Network Topology

A brain network is a weighted graph **g** = (*V, E*) consisting of a node set 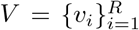, where *v*_*i*_ is a brain region out of *R* total regions, and edge set *E* = {(*v*_*i*_, *v*_*j*_)|*i ≠j*} whose weights *A*_*ij*_ = *a*(*v*_*i*_, *v*_*j*_) ∈ [0, 1] quantify the strength of the connections between each pair of nodes. In functional connectivity studies, *v*_*i*_ are determined based on a parcellation of brain regions and *A*_*ij*_ are often calculated using the absolute value of the between-region correlations of fMRI time series (Achard, 2006). The adjacency matrix associated with **g** is then defined as the matrix *A* with *i, j*th entry equal to *A*_*ij*_.

The topology of **g** can be described using network diagnostics (Bullmore and Bassett, 2011) that are lower dimensional than *A*, but that are more informative about the overall network organization than a single edge. For a graph **g** and any partition of its nodes into communities, the modularity of **g** describes the abundance of connections within communities that we observe compared to a null graph:

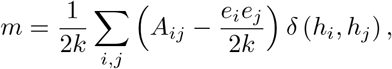

where *e*_*i*_ is the degree of (i.e., the number of edges connecting to) node *i, k* is the total number of edges, *h*_*i*_ is the community assigned to node *i*, and *δ* is the Kronecker delta. Intuitively, when nodes *v*_*i*_ and *v*_*j*_ are in the same community, the term 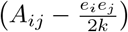 is the difference between the observed connectivity *A*_*ij*_ and 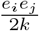, the expected connectivity of a random graph under a configuration null model (Newman and Girvan, 2004). In the configuration null, the degree of each node is preserved, but all *k* edges are cut in half, creating 2*k* “open” half-edges. Then, the network is re-wired: all 2*k* open half-edges are randomly connected to each other with equal probability, a process that allows for self-loops and multiple edges between any two nodes. Therefore, any edge emanating from a node has 2*k* − 1 possible locations to connect to. For two nodes *v*_*i*_ and *v*_*j*_ with degree *e*_*i*_ and *e*_*j*_, respectively, the probability of an edge existing between them is 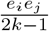, which is often approximated by 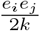 when *k* is large. A larger value of 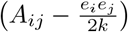, and hence modularity, means that more connections within communities are observed than might be expected under the configuration null. Community assignments *h*_*i*_ may either be known *a priori* or calculated as part of an iterative and greedy modularity maximization algorithm (Blondel *and others*, 2008; Fortunato, 2010). In our paper, we use pre-defined communities based on a 7-community node partition resulting in functionally coupled clusters in the cerebral cortex; these are described in further detail in Yeo *and others* (2011).

Similarly, the participation coefficient (Guimerà and Amaral, 2005) captures how often the average node connects to nodes outside its own community, and is calculated as

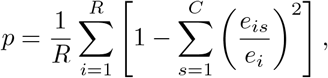

where *e*_*is*_ is the number of edges between node *v*_*i*_ and all nodes in community *s, e*_*i*_ is the degree of node *i*, and *C* is the total number of communities. When the term 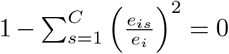, node *i* is only connected to nodes in its own community; when it equals 1, node *i* is equally connected to every community.

### 2.2. Network Thresholding and Diagnostic Curves

Thresholding is a common technique when the goal is to produce binary networks, or weighted networks where the weakest edges—lower correlation values that are most impacted by measurement error—have been removed (Zalesky *and others*, 2012; Bordier *and others*, 2017). Because two networks with different numbers of non-zero edges may not be comparable in terms of their network diagnostics (Garrison *and others*, 2015; van Wijk *and others*, 2010), we threshold the edges such that the same proportion of strongest edges is preserved. The fraction *t* of edges preserved is the proportional threshold (Geerligs *and others*, 2014; Bassett *and others*, 2012).

As the choice of threshold *t* is often arbitrary, we instead consider network diagnostics as a function of all *t* ∈ [0, 1]. For person *i* = 1, …, *n* and threshold value *t*_*k*_ for *k* = 1, …, *r*, we observe a single weighted network upon which we can calculate the modularity *m*_*i*_(*t*_*k*_) and participation coefficient *p*_*i*_(*t*_*k*_) (Figure 1). By repeating this process for all *i* and *t*_*k*_, we obtain the discrete observations 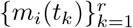 and 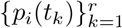, which are then assumed to be samples of the smooth curves *m*_*i*_(*t*) ∈ *ℒ*^2^[0, 1] and *p*_*i*_(*t*) ∈ *ℒ*^2^[0, 1].

Functional data analysis methods (Wang *and others*, 2016) automatically encode the autocorrelation of the network diagnostic between neighboring values of *t*. Because functions are infinite-dimensional objects (i.e., they can be parameterized by infinitely many parameters), they are often replaced with a lower-dimensional representation during model estimation. One widely-used approach is functional principal components analysis (FPCA), which can simultaneously perform dimension reduction and decompose the functional variance in a set of curves. Letting *μ*_*m*_(*t*) and Σ_*m*_(*s, t*) ≡ Cov(*m*_*i*_(*s*), *m*_*i*_(*t*)) be the mean and covariance functions of *m*_*i*_(*t*), we can express *m*_*i*_(*t*) as

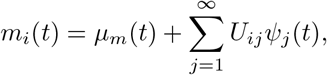

where *ψ*_*j*_(*t*) is the *j*th eigenfunction or functional principal component (FPC) of Σ_*m*_(*s, t*) based on its spectral decomposition, and *U*_*ij*_ is the associated FPC score

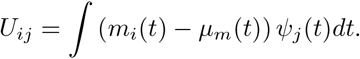

Typically, this infinite sum can be well approximated by the first *N*_*m*_ terms (Wang *and others*, 2016):

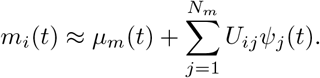

In practice, the observed curves *m*_*i*_(*t*) may be conceptualized as true curves 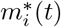 distorted by measurement error:

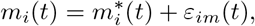

where *ε*_*im*_(*t*) is a mean zero measurement error process with variance 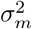. Then, Σ_*m*\*_ (*s, t*) can be estimated by smoothing the method of moments estimator of Σ_*m*_(*s, t*) from the observed data (Goldsmith *and others*, 2011). Putting these together, we can write the FPCA expansion of *m*_*i*_(*t*) as the sum of the mean, a linear combination of finitely many FPCs, and measurement error:

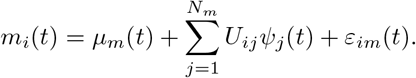

Similarly, we write the FPCA decomposition of *p*_*i*_(*t*) as

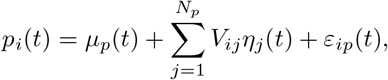

allowing us to express the participation coefficient curves again as the sum of the mean, a linear combination of FPCs, and measurement error.

### 2.3. Scalar-on-Function Regression

To relate a scalar-valued outcome *y*_*i*_ to a function-valued covariate *m*_*i*_(*t*), we first consider the simplest possible scalar-on-function regression model (Ramsay and Silverman, 2005):

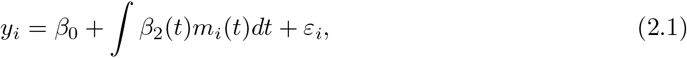

where *ε*_*i*_ ∼ *N* (0, *σ*^2^) is independent Gaussian noise, *β*_2_(*t*) is a smooth coefficient function, and *β*_0_ is an intercept. Model 2.1 can be estimated assuming that *m*_*i*_(*t*) is replaced with its low-dimensional FPCA decomposition, and *β*_2_(*t*) can be represented in terms of penalized splines. Thus, a data-driven way to identify the thresholds *t* where *m*_*i*_(*t*) influences the outcome *y* is to find the values of *t* for which *β*_2_(*t*) is substantially different from zero.

### 2.4. Curve Alignment

Proportional thresholding can introduce a new type of confounding if there are systematic differences in the distribution of network edge weights (van den Heuvel *and others*, 2017). For example, brain networks in older children tend to have stronger connections overall, compared to younger children. As a result, the strongest 15% of edges for older participants are less likely to contain spurious connections (which have a more random topology) than the same percentage of edges in younger participants. This fact implies that for persons *i≠ j, m*_*i*_(*t*_0_) and *m*_*j*_(*t*_0_) may not be comparable at the same value *t*_0_.

When the modularity curves *m*(*t*) are functional objects, this lack of comparability can be conceptualized as phase (horizontal) variation: although the curves *m*_*i*_(*·*) and *m*_*j*_(*·*) describe the same underlying process, one curve may be a horizontally shifted or more intricately warped version of the other (Figure 2). Inference based on amplitude (vertical) variation—the difference in the height of the modularity curve at fixed values of *t*—may be biased in the presence of phase variation, since phase variation precludes the accurate calculation of fundamental quantities such as the mean curve. As a result, methods for functional alignment (also called curve registration or warping) have been extensively studied and reviewed (Marron *and others*, 2015; Wang *and others*, 2016; Wrobel *and others*, 2018; Srivastava *and others*, 2011). Once they are aligned to a new functional domain, the amplitude variation of modularity and participation curves (e.g., the difference between *m*_*i*_(*t*_0_) and *m*_*j*_(*t*_0_) for persons *i* and *j* and time *t*_0_) can be meaningfully interpreted as differences in network structure. A common strategy is to address phase and amplitude variation separately: functions are pre-aligned before applying a downstream analysis such as regression or clustering (Reiss *and others*, 2017).

**Fig. 2.**
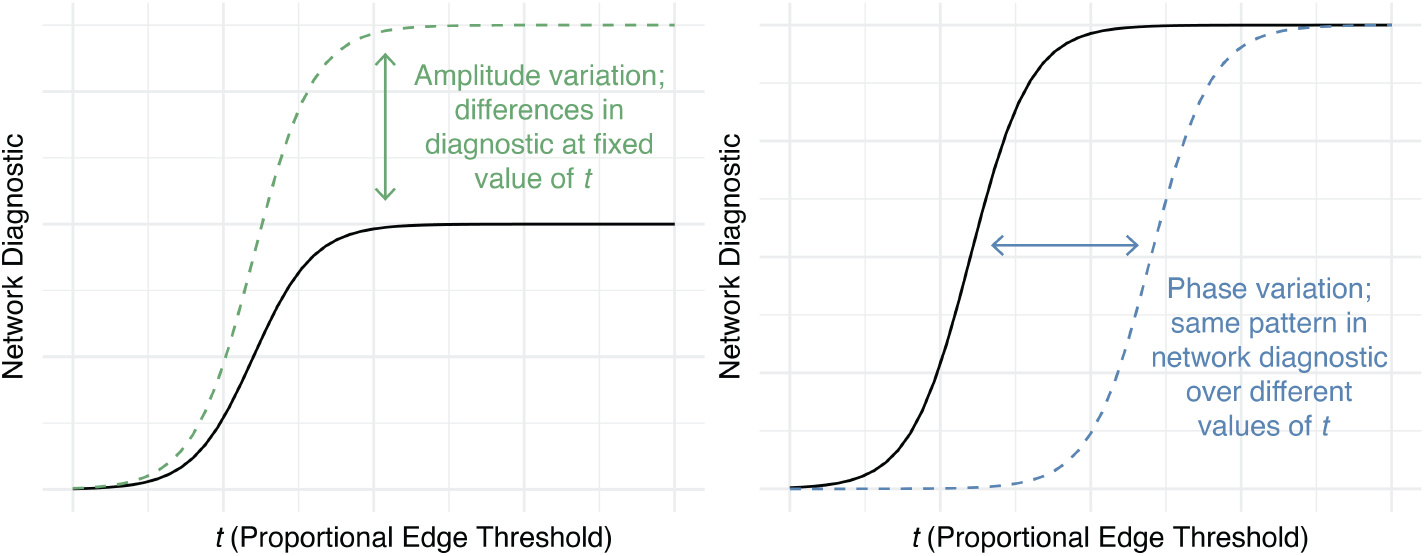
Forms of variation in functional data. Left: For network diagnostic curves that are functions of *t*, amplitude variation refers to differences in the vertical distance between curves at fixed values of *t*. Right: For curves with similar features, phase variation manifests as differences in the “timing” of those features, i.e., horizontal misalignment.

An important note is that curve alignment and scalar-on-function regression are two methods that frequently occur in tandem but differ in their ends, given the structure of our data. Since the network diagnostic curves are simple and monotonic, it is possible to perfectly align them. But if the curves are perfectly aligned such that *m*_*i*_(*t*) = *m*(*t*) for all *t*, then Model 2.1 cannot be estimated. In general, the more these curves overlap (i.e., the less variation between curves), the more unstable the estimate of *β*_2_(*t*) is likely to be. In other words, we need variation in the predictors to draw any conclusions about associations with the response. At the same time, estimating Model 2.1 without performing any warping of the data may lead to incorrect conclusions, as stated above. We therefore investigate if it is possible to find an optimal middle ground between no alignment and perfect alignment in the case of simple, monotonic curves. Note that this issue is less of a concern when working with other types of functional data that have multiple non-monotonic features and residual differences in amplitude that remain after alignment.

Another concern is that, due to the flexibility of the scalar-on-function regression model, it is likely that there will be multiple ways to warp the data such that the predicted value of *y* is exactly equal to the observed value (i.e., the model is overfit to the data). Our proposed method aims to warp the data such that the prediction of an *auxiliary* variable *z* is improved, and we explore whether alignment using *z* can improve the prediction of *y* later on, regardless of the relationship between *y* and *z*.

Previous methods have considered the joint estimation of regression parameters and warping functions in the context of function-on-scalar and function-on-function regression. In Gervini (2014), the amplitude variation is decomposed using FPCA and the phase variation decomposed using Hermite splines; the associated scores and knots of these decompositions are then related to scalar covariates through a linear mixed effects model. However, as noted in Tucker *and others* (2018), this method involves applying FPCA to the unaligned curves, which is suboptimal as much amplitude variation may be artifactual due to misalignment. In Hadjipantelis *and others* (2015), a function-on-scalar model is proposed where both the amplitude and phase variation are decomposed using FPCA and the associated FPC scores are related to covariates. In Tucker *and others* (2018), phase and amplitude variation are simultaneously assessed via a joint FPCA, and then the outcome is regressed on these FPCs. However, FPCA is performed in a transformed space via the square root velocity function, which renders the principal components less interpretable. Our proposed model differs from these in that we consider a scalar-on-function framework where the outcome *y* is regressed onto the aligned curves *m*_*i*_(*t*), which are aligned in a supervised way, involving an auxiliary variable *z* that is related to the aligned *m*_*i*_(*t*) and possibly *y*. The proposed alignment algorithm alternates between estimation of the parameters of a model for *z* given the warped curves, and the warping functions given the *z* model.

### 2.5. Warping Functions

As others have done, we differentiate between the “individual threshold” 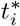 of the observed unaligned curves, and the “system threshold” *t* of the aligned curves (Wrobel *and others*, 2018), which are connected through the warping function 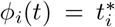. Therefore, the model in (2.1) becomes

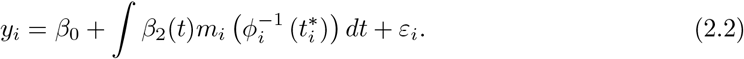

In the new functional domain, regions of *t* where the coefficient function *β*_2_(*t*)*≠* 0 corresponds to the portions of the *aligned* curves that are most associated with the outcome. In other words, *β*_2_(*t*) encodes information about the relationship that exists between the outcome and network curves without the effects of phase variability due to age. Note that Model (2.2) involves the aligned curves 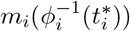, though we only observe the unaligned curves 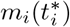.

As network diagnostic curves tend to be simple in shape, we choose a simple parameterization for the inverse warping functions,

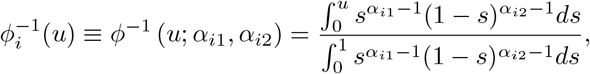

the regularized incomplete Beta function with parameters *α*_*i*1_, *α*_*i*2_ *>* 0. The function 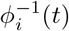 is the cumulative distribution function (CDF) of a Beta(*α*_*i*1_, *α*_*i*2_) random variable, and is naturally monotonic and constrained at the endpoints so that *ϕ*^−1^(0; *α*_*i*1_, *α*_*i*2_) = 0 and *ϕ*^−1^(1; *α*_*i*1_, *α*_*i*2_) = 1 for all *α*_*i*1_, *α*_*i*2_ *>* 0. For identifiability, alignment methods often require that the warping function is equal to the identity on average: 𝔼 [*ϕ*(*t*)] = *t*, though many prefer to enforce 𝔼 [*ϕ*^−1^](*t*) = *t* for computational convenience (Fu and Heckman, 2019). Here, we note an advantage of using a CDF-based warping function: both *ϕ*^−1^(*t*) and its inverse *ϕ*(*t*)—a quantile function—are simple to evaluate numerically. The requirement that 𝔼 *ϕ*^−1^(*t*) = *t* can also be replaced with the weaker requirement that 𝔼 [(*α*_*i*1_, *α*_*i*2_)] = (1, 1).

Another advantage of selecting warping functions with a closed-form inverse is the capacity to plan simulation studies. In the simulations presented in this paper, the inverse warping function (e.g., the Beta quantile function) can be used to warp the data using known parameters that allow for the recovery of the accurately aligned curves with the original warping function (e.g., Beta CDF). Subsequently, these parameters can be compared with the estimated parameters derived from our algorithm. However, there are few well-explored classes of warping functions that are both continuous and bijective on [0, 1] and possess a closed-form inverse. While our paper primarily focuses on the Beta CDF, we will also gauge alignment performance using an alternative class of simple polynomial warping functions *ξ*(*t*) = *t*^*α*^, where *t* ∈ [0.1]. Other choices of warping functions satisfying the inverse property include piece-wise linear warping functions (McDonnell *and others*, 2021), which requires constrained optimization over four parameters, and CDFs of other truncated distributions.

Even with the aforementioned restrictions, Model (2.2) is not identifiable without further assumptions, since the true aligned curves are not observed for any person. A full likelihood involving the penalized regression parameters and fixed effects *α*_*i*1_, *α*_*i*2_ would be over-parameterized, with the number of parameters increasing with the number of participants *n*. Therefore, we consider an iterative and penalized likelihood heuristic to estimate *α*_*i*1_, *α*_*i*2_ given an auxiliary variable *z*, with the ultimate goal of using the aligned functions to predict an outcome *y*. The reason we do not use *y* itself for alignment is that, in many settings, it is possible to find *α*_*i*1_, *α*_*i*2_ that results in a perfect fit (i.e., *y*_*i*_ = *ŷ*_*i*_) even when there is no true association between *y* and the predictor curves. Therefore, we should believe *a priori* that the auxiliary variable *z* is strongly associated with the functional covariate (as we do with age and the network characteristics discussed above).

### 2.6. sRAFT Algorithm

The RAFT algorithm estimates Model (2.2) using a two-step iterative process. In the model step, we estimate the parameters for the model involving *z* and *m*_*i*_(*t*); for instance, scalar-on-function regression with coefficients *θ* ≡ *{β*_*z*0_, *β*_1_(*t*)}. In the warp step, we estimate warping functions *ϕ*_*i*_(*t*) and aligned curves 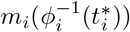.

1. **Initialize** at no warping: 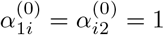 for *i* = 1, *· · ·, n*.
2. **Model step**. At iteration *k*, fit a penalized functional regression model for outcome *z* with parameters *θ*^(*k*)^ given 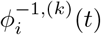 and the warped data 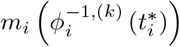:

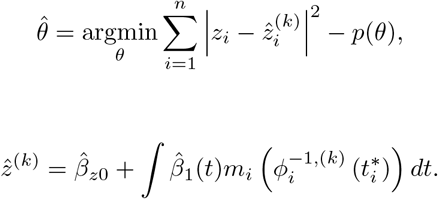

The functional coefficient *β*_1_(*t*) is estimated using penalized B-splines with a large number of knots 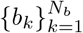 with a smoothness-inducing penalty term *p*(*θ*) (Goldsmith *and others*, 2011). The functional predictor is represented using its FPCA decomposition:

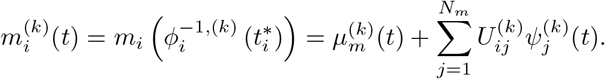

The model above can be readily estimated using scalar-on-function regression software, from the `refund R` package (Goldsmith *and others*, 2011, 2016).
3. **Warp step**. Estimate warping functions 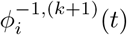 given *θ*^(*k*)^:

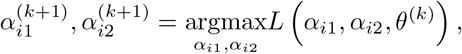

where the objective function

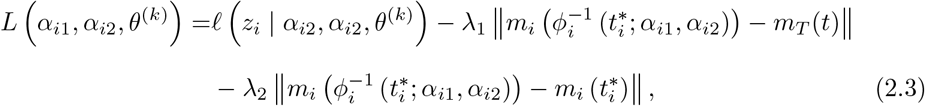

and the term ℓ(*z*_*i*_ | *α*_*i*2_, *α*_*i*2_, *θ*^(*k*)^) is the likelihood of *z* assumed in Step 2.
4. **Iterate** steps 2 and 3 until convergence: 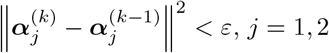.
5. Finally, we fit the scalar-on-function model for the outcome of interest *y*,

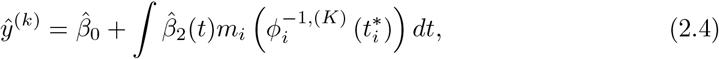

where 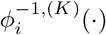 is the final warping function after *K* iterations.

The objective function in Step 3 has three terms. Intuitively, the warp 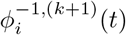 is chosen to maximize the likelihood

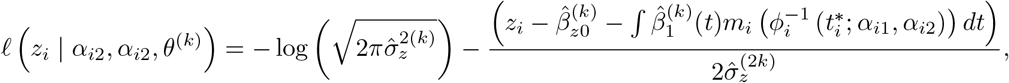

given the observations *z*_i_ and estimated coefficients 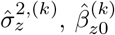, and 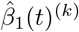, but within certain parameters: the second term in *L* (*α*_*i*1_, *α*_*i*2_, *θ*^(*k*)^)ensures that the data are warped towards a template curve, which can be known *a priori* or selected in a data-driven manner. For example, we select the centroid curve in the data which minimizes the *L*^2^ distance from all other curves. The third term ensures that the warped curve does not deviate too far from the original curve. Together, the second and third terms of *L* (*α*_*i*1_, *α*_*i*2_, *θ*^(*k*)^) encourage a balance between alignment towards the template and maintaining the original curve. We propose that an *unsupervised* warping procedure can be done in a single step with the modified loss function

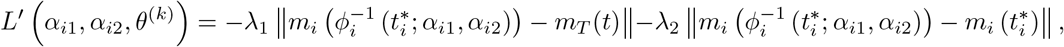

where the likelihood term—the only term involving the supervising variable *z*—is removed.

#### 2.6.1 Tuning Parameters

The parameters *λ*_1_ and *λ*_2_ control the balance of these penalties: when *λ*_2_ is very large, no warping is performed, and when *λ*_1_ is very large, curves are aligned towards the template. The values of these parameters must be chosen ahead of time or through an automated procedure. In the latter case, we suggest setting aside a portion of the data as a tuning set, where the data are warped using *λ*_1_ and *λ*_2_ selected from a grid of possible values, and the optimal *λ*_1_ and *λ*_2_ are those which maximize the R-squared of Model (2.4). Having selected the tuning parameters (separately for supervised and unsupervised warping), the algorithm is then applied to the remainder of the data.

#### 2.6.2 Generalized Algorithms

In principle, the model for *z* in Step 2 of the RAFT algorithm can be any method for scalar-on-function regression, such as non-parametric regression or a suitable machine learning algorithm. In that case, the term ℓ (*z*_*i*_ | *α*_*i*2_, *α*_*i*2_, *θ*^(*k*)^)in the objective function may be replaced by another criterion, such as the MSE. Finally, in addition to the “model-first” algorithm above, where the model for *z* is estimated after initialization and before the first iteration of warping, it is also possible to consider a “warp-first” algorithm where the initial warps are estimated using an unsupervised step initially.

## 3. Simulation Results

Performance of RAFT versus similar methods was first gauged using simulated data under multiple conditions. We first describe the simulation strategy at a high level; further details about the specific data generating process can be found in Section 3.1 below. The estimated penalty parameters are listed in Table S.1.

We generated *n* = 600 curves with the same base curve shape *f*_0_(*t*) for all individuals, similar to those used by Carroll *and others* (2020). Two strategies were used to induce amplitude (vertical) variation between these base curves: sinusoidal noise, where a random sine and cosine curve defined on [0,1] were added to *f*_0_(*t*) (Figure 3, Panels A and C); and Gaussian noise, where a random Gaussian variable was added to *f*_0_(*t*) in a pointwise fashion (Figure 3, Panels B and D). Finally, we considered two coefficient functions *β*_1_(*t*) and *β*_2_(*t*) representing the true association between the aligned curves and (*z, y*). Thus, the true data generating process involved the aligned curves *f*_*i*_(*t*):

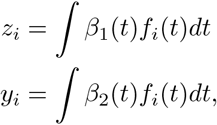

while the observed data include the unaligned curves *f*_*i*_(*ϕ*(*t*; *α*_*i*1_, *α*_*i*1_)) and (*z, y*), with the goal to accurately estimate *β*_2_(*t*), as *y* is the ultimate outcome of interest.

**Fig. 3.**
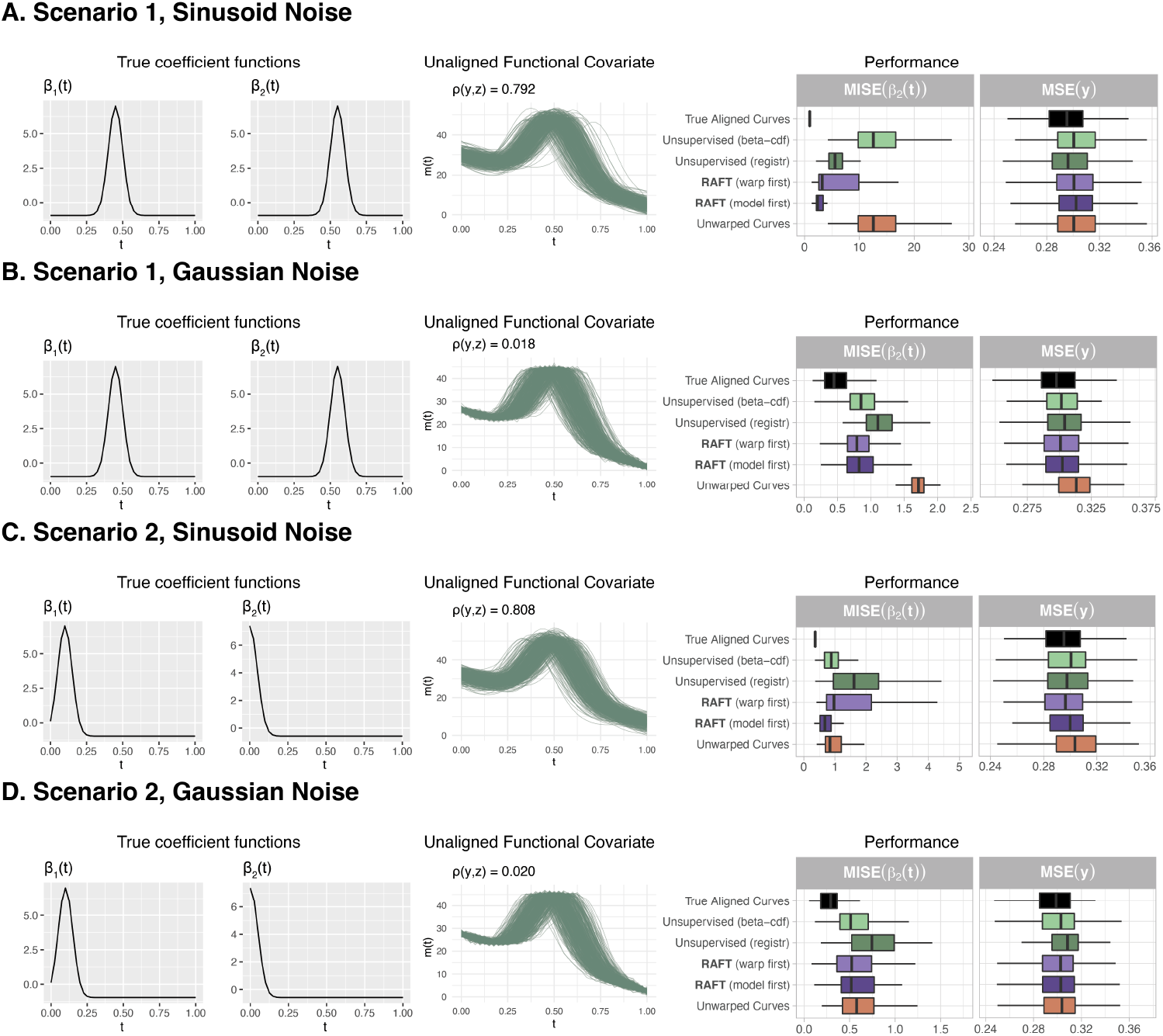
In simulated data, alignment improves the accuracy of model coefficient estimation beyond using the unwarped curves, and towards the scenario where the true curves are known. The data generating mechanism consisted of a set of true *β*_1_(*t*) and *β*_2_(*t*) functions describing the association between the true aligned curves and *z* and *y*, respectively, where sinusoidal noise (Panel A) and Gaussian noise (Panel B) were added to the functions to induce amplitude variation. This process was repeated for curves with added sinusoidal noise (Panel C) and Gaussian noise (Panel D) under a different set of true *β*_1_(*t*) and *β*_2_(*t*) functions. For the supervised alignment methods, the tuning parameters *λ*_1_, *λ*_2_ were chosen in that order, and using the automated method in 2.6.1. Performance was assessed using MISE(*β*_2_(*t*)) and MSE(*y*). Box-and-whisker plots show the median, 25th, and 75th percentile for the box, and 1.5 * (Interquartile Range) for the whiskers. Overall, we found that RAFT most consistently improved prediction performance, even when the variables *z* and *y* were not linearly related (Panels B and D), while the unsupervised FPCA-based method *registr* worked often but occasionally performed worse than using the unwarped curves, likely due to the increased flexibility in modeling the warping functions.

Model performance was measured by the mean squared error (MSE) of *y* and mean integrated squared error (MISE) of *β*_2_(*t*) over 100 iterations. For the supervised methods, the tuning parameters (*λ*_1_, *λ*_2_) were automatically selected as described in Section 2.6.1 on a tuning set of *n* = 125 other curves, and performance was assessed on the remaining 500 curves. In addition to the proposed methods from this paper—model-first RAFT, warp-first RAFT, and unsupervised “Beta CDF” warping—we also compared our results to *registr*, an FPCA-based unsupervised warping method (Wrobel *and others*, 2018; Wrobel, 2018). These four alignment methods were evaluated against the performance levels reached when using either the unaligned curves and the true aligned curves.

We found that aligning the functional covariate almost always improved (lowered) MISE(*β*_2_(*t*)) beyond simply using the unaligned curves to predict *y* (Figure 3). In scenario 1, model-first and warp-first RAFT had the best MISE(*β*_2_(*t*)), closest to that observed in the ideal case where the true aligned curves are known. In scenario 2, RAFT methods performed well again, but the unsupervised Beta CDF alignment proposed in Section 2.6.1 also performed well. The true coefficient *β*_2_(*t*) in scenario 2 has a sharp increase at *t* = 0, suggesting that for this type of relationship, FPCA-based warping methods such as *registr* may be less optimal than the more constrained warping provided by our proposed Beta CDF alignment. The value of in-sample MSE(*y*) did not vary appreciably between any of these methods.

Inferential properties of each method were assessed in terms of the power and coverage probabilities of the pointwise confidence bands of 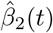(Marra and Wood, 2012). In general, power and coverage probability were highest in the ideal case where the true aligned curves were known, and lowest without any alignment, with the alignment methods performing moderately (Figure S.1). Among the alignment methods, *registr* had best power and coverage probabilities when the true *β*_2_(*t*) was further from zero and in the presence of sinusoidal noise in the covariate functions. In the presence of Gaussian noise, both RAFT supervised and unsupervised alignment exhibited power and coverage that were closest to the ideal case.

### 3.1 Simulation Details

The base curve had the form 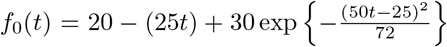. To add sinusoidal noise to the base curve, we generated observed curves *f*_*i*_(*t*) = *f*_0_(*t*) + *u*_1*i*_ sin(2*πt*) + *u*_2*i*_ cos(2*πt*), with i.i.d. *u*_*ji*_ ∼ *N* (*μ* = 0, *σ* = 0.5*r*) and *r* = max |*f*_0_(*t*)) − min(*f*_0_(*t*)|. To add Gaussian noise to the base curve, we generated observed curves *f*_*i*_(*t*) = *f*_0_(*t*) + *u*_*i*_(*t*) with i.i.d *u*_*i*_(*t*) ∼ *N* (*μ* = 0, *σ* = 0.015*r*) independently for all *t*. The final sample of observed but unaligned curves were then generated as 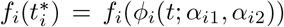, with i.i.d. 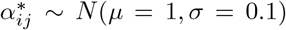 for *j* = 1, 2 and 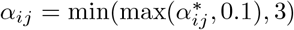 to ensure all *α*_*ij*_ *>* 0. We found that values of *α*_*ij*_ *>* 3 and *α*_*ij*_ *<* 0.1 produced extreme warps, where the entire curve was compressed towards one of the endpoints.

We also considered two options for the true coefficient functions *β*_1_(*t*) and *β*_2_(*t*). In case 1, 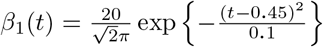 and 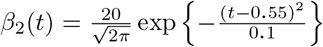 for *t* ∈ [0, 1]. In case 2, 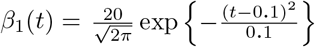 and 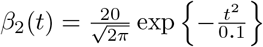. Both sets of coefficient functions were chosen to be more hill-shaped so that results were more sensitive to the effects of misalignment.

Pointwise confidence bands around 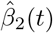 were obtained by calculating the standard error (uncorrected for bias) multiplied by two, as implemented in the *mgcv* and *refund* R packages (Marra and Wood, 2012; Goldsmith *and others*, 2016). For each value of *t*, power was calculated as the proportion, over 100 repetitions, where the confidence band excluded 0 if *β*_2_(*t*)*≠* 0. (In the simulations, *β*_2_(*t*) was non-zero at all evaluation points *t* ∈ {0, 0.01, …, 1}.) Coverage probability was computed as the proportion of repetitions where the confidence band included the true value of *β*_2_(*t*).

### 3.2. Polynomial Warping Functions

In addition to the Beta CDF warping functions introduced in this paper, we also experimented with polynomial warping functions *ξ*(*t*) = *t*^*α*^ (discussed in Section 2.5) to align the curves. Although the polynomial warping function is less flexible than the Beta CDF function in that it does not have any inflection points (i.e., where the curvature can change sign), the polynomial function depends on only one parameter instead of two, which may improve algorithm performance. We repeated the same experiment outlined in Section 3.1 and found that the polynomial warping functions performed similarly to the Beta CDF warping functions in most scenarios, but impacted the unsupervised method (resulting in lower performance) and the warp-first RAFT method (resulting in better performance) (Figure S.2).

### 3.3. Sensitivity to Template Choice

We evaluated how the choice of the template function could affect the performance of the supervised and unsupervised algorithms. While our method uses the *L*^2^ centroid as the template by default, in sensitivity analyses, we also considered simply setting the template as a random observation from the data. Overall, the alignment methods suffered a moderate drop in performance for Scenario 1 and performed similarly in Scenario 2 (Figure S.4).

We further considered the situation where the template was chosen as the default *L*^2^ centroid, but was warped again by a known amount using a *ϕ*(*t*, 0.5, 1.5) warping function so that it was less representative of the data. In this setting, the supervised and unsupervised methods all worsened in terms of MISE(*β*(*t*)), often performing worse than any other competing method (Figure S.5).

### 3.4. Sensitivity to Covariate Warping and Shape

In the final set of sensitivity analyses, we experimented with alternative data generating mechanisms. We replicated our analyses in Section 3.1, but for base curves of the form 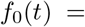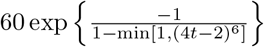, which have a bump-shaped plateau (Figure S.3) resembling those of 24-hour activity profiles (Leroux *and others*, 2019). Mirroring our main results, the RAFT and unsupervised methods performed well compared to no alignment, though *registr* had equal or better performance in this setting. Notably, in Scenario 1, the estimated *λ*_2_ values were high when sinusoidal noise was added to the data, which resulted in no warping for either the supervised or unsupervised methods.

We also checked the assumption that our method should behave similarly if all of the unaligned curves were uniformly warped (meaning that the same warping function was applied to the unaligned covariate functions). In simulations, we applied an additional *ϕ*(*t*, 0.75, 0.75) warping to each of the unaligned curves prior to alignment. In contrast to the case when only the template was warped, a uniform warping of all the covariates either slightly improved performance or left it unchanged (Figure S.6).

## 4. Community Structure and Cognitive Performance in Neurodevelopment

We next asked if modularity and participation coefficient curves could improve prediction of executive function, a *z*-scored quantity representing non-verbal reasoning ability (Appendix S.1), once the curves had been warped using either age or motion information. To do so, we applied our method to *n* = 1122 participants from the Philadelphia Neurodevelopmental Cohort (Satterthwaite *and others*, 2014) who underwent functional connectivity and neurocognitive studies. For each participant, the functional connectivity matrix was calculated by taking the correlation between the averaged BOLD time series in 400 cortical regions defined by Schaefer *and others* (2017). To generate a sequence of thresholded weighted matrices, the correlation matrix was cumulatively thresholded by a proportional threshold value *t* = {0.01, 0.02, …, 0.99, 1}. For each person and value of *t*, we obtained a weighted network on which we calculated the modularity and participation coefficient values given the 7-community node partition described in Yeo *and others* (2011).

We also considered scalar covariates that are often studied in conjunction with functional connectivity in youth: age in years and in-scanner subject motion measured in millimeters (mm). Motion, defined as the average relative root mean square (RMS) displacement, has been shown to impact functional connectivity estimates and downstream network diagnostics (Power *and others*, 2012). Our data were pre-processed to reduce motion effects and all participants had average RMS displacement *<* 0.2 mm per frame as part of the inclusion criteria, although residual motion effects may persist (Ciric *and others*, 2017; Satterthwaite *and others*, 2014). Further details on data collection and scanner settings can be found in Appendix S.1.

Older adolescents had less motion (Figure S.8) and stronger, heavier-tailed distributions of raw functional connectivity (FC) edge weights, i.e. a greater proportion of strong connections (Figure 4). Specifically, we observed weak positive correlations (*ρ*, [95% CI]) between age and the following: average edge weight, *ρ*=0.197 [0.140,0.254]; standard deviation of edge weights, *ρ*=0.217 [0.161,0.272]; and the inter-quartile range of edge weights, *ρ*=0.205 [0.149,0.261]. We found weak negative correlations between motion and the following: average edge weight, *ρ*=-0.155 [-0.097,-0.211]; standard deviation of edge weights, *ρ*=-0.191 [-0.247,-0.134]; and the inter-quartile range of edge weights, *ρ*=-0.173 [-0.230,-0.116].

**Fig. 4.**
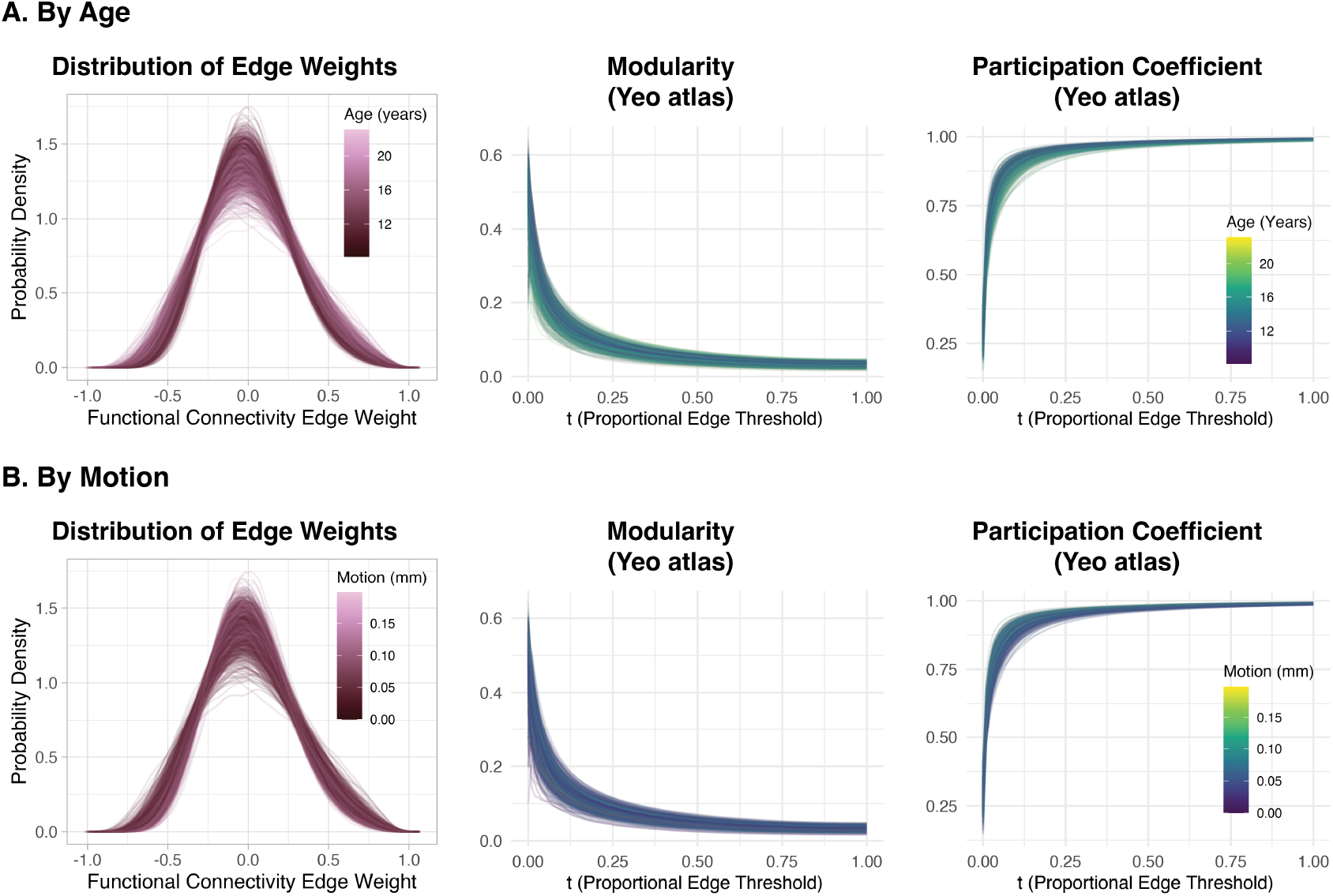
Data from the Philadelphia Neurodevelopmental Cohort (Satterthwaite *and others*, 2016). Left column: The distribution of functional connectivity (FC) strength, which determine network edge weights, is related to age (Panel A) but not motion (Panel B). Middle and right columns: Modularity and participation coefficient given the 7 communities defined in Yeo *and others* (2011) as a function of *t*. The color of the curve corresponds to age (Panel A; darker = younger) or motion (Panel B; darker = less motion).

Figure 4 shows the network diagnostic curves in our sample. Modularity curves followed a similar shape, with a sharp peak around *t* ≈ 0.05 and then a downward slope as *t* increased. There was greater amplitude variation of the curves when *t <* 0.5. Participation coefficient curves followed a complementary path, with an upwards-sloping shape as *t* increased: they then levelled off at around *t* = 0.4 and remained relatively flat thereafter.

We then compared various alignment methods, including no alignment, in terms of their ability to predict executive function. This process involved fitting Model (2.2), where the outcome *y* was executive function and the covariate 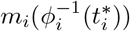 was the aligned participation coefficient or modularity curve. The estimated penalty parameters are shown in Table S.2. As in the simulations, we compared supervised warping (model-first and warp-first RAFT) with *registr* and unsupervised warping methods. Because the true aligned curves and model coefficients are not observed, the performance of the alignment methods was gauged using the MSE of *y* and the corresponding R-squared.

Supervised alignment was performed using either the auxiliary variable *z* = age (years) or *z* = motion (mm), as both are thought to be related to observed FC. As before, the tuning parameters (*λ*_1_, *λ*_2_) in the supervised alignment method were selected using the method in Section 2.6.1. The data were split into 5 folds, and for each iteration (out of 5), one fold was the tuning set and the other 4 folds were the testing set on which the MSE(*y*) and R-squared were evaluated. This process ensured that any data used to tune (*λ*_1_, *λ*_2_) in one iteration could later be re-used as testing data.

When the auxiliary warping variable *z* is age, then the curve alignment with RAFT resulted in a model with lowest MSE(*y*) and consequently highest R-squared (Figure 5). By comparison, unsupervised methods did not significantly improve performance beyond that of the unaligned curves. This behavior was observed for both the modularity and participation coefficient curves. When the auxiliary warping variable *z* is motion, then no method outperformed the others in predicting executive function, in that they each had a high MSE(*y*) and R-squared approximately equal to 0. In other words, alignment using motion did not improve model performance beyond no alignment whatsoever, and in either case the model fit was poor.

**Fig. 5.**
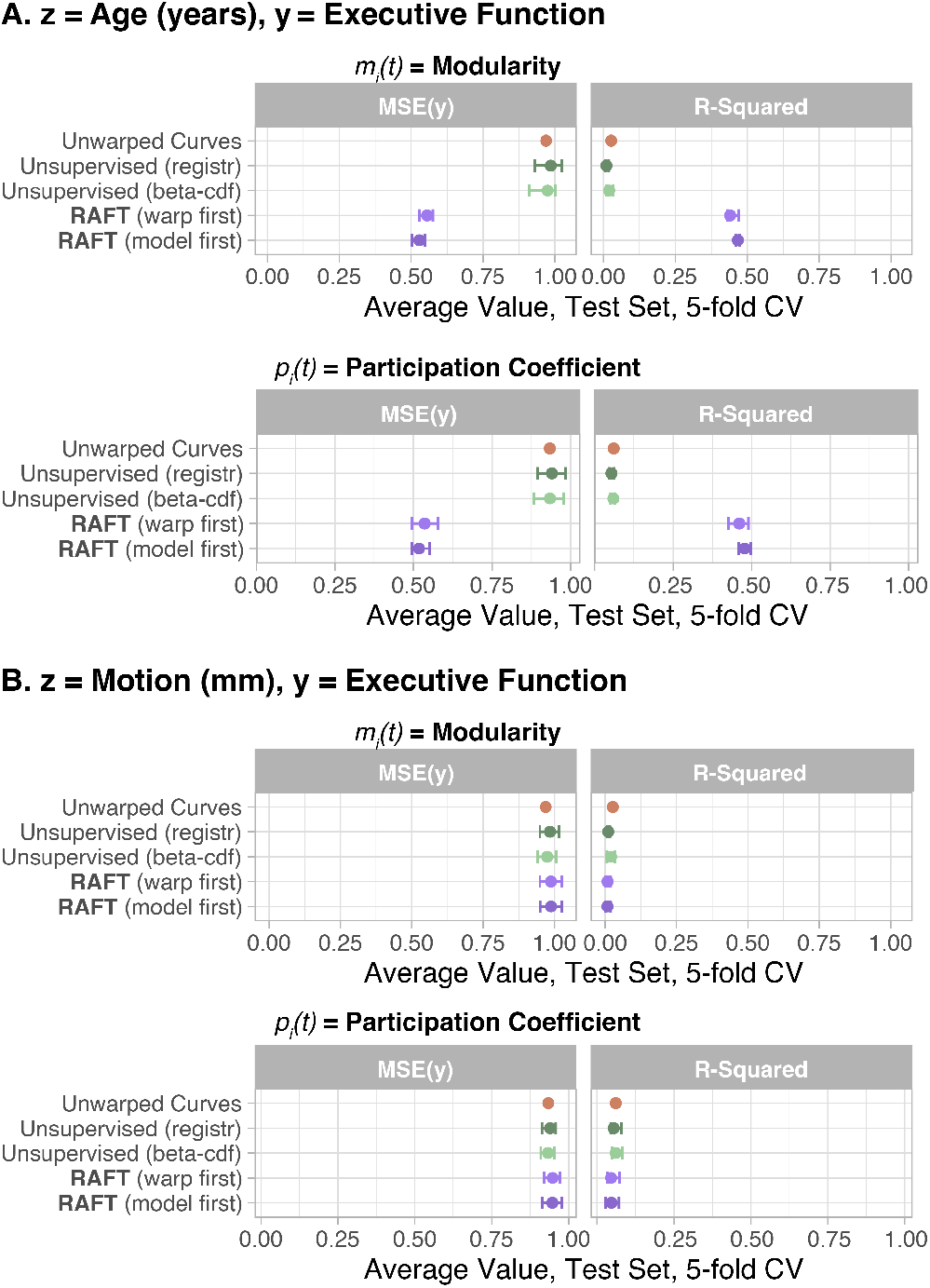
In data from the Philadelphia Neurodevelopmental Cohort (Satterthwaite *and others*, 2016), RAFT alignment of the modularity and participation coefficient curves improved prediction of executive function when the auxiliary variable was age, but not when the auxiliary variable was motion. We considered modularity (left column) and participation coefficient (right column) using communities defined by the 7-community parcellation described in (Yeo *and others*, 2011); age was measured in years, and motion measured as average relative RMS displacement in millimeters. (Panel A) The superior performance of RAFT when *z* is age could be due to multiple factors: age has a strong association with both the network diagnostic curves (Figure S.13 and Figure S.14) and executive function. Therefore, warping the curves to predict age well will likely improve prediction of *y*. Another possibility is that age-related misalignment has obscured the true association between executive function and network diagnostic curves, which RAFT has recovered. (Panel B) The similarly poor performance of all alignment methods, including RAFT, when *z* is motion could be due to the weak association between *z* and the network diagnostics, leading to a warping that look similar to the unsupervised case.

We also asked if the functional predictors provide information beyond *z* by including the latter in the regression model for predicting *y*. When a linear term for *z*_*i*_ was included in the model so that

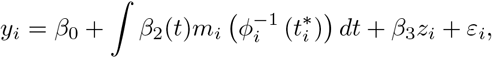

then the RAFT alignment still out-performed other methods for when *z* is age but not for when *z* is motion (Figure S.9). However, when a non-linear smooth term *s*(*z*_*i*_) using penalized regression splines was included in the model so that

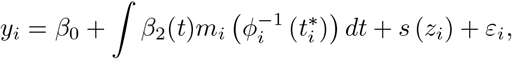

then the RAFT alignment did not result in performance improvements (Figure S.10). In other words, there was similar performance when predicting executive function using either solely the curves warped using age, or the combination of the aligned curves and the highly flexible smooth age term *s*(*z*_*i*_).

By examining the estimated coefficient functions (Figure S.15), we found an association between executive function and lower modularity (after alignment) for lower thresholds, similar to that observed in the unaligned curves. However, supervised alignment methods also revealed significant associations at other regions of the modularity curves; in contrast, *registr* -aligned modularity curves were not associated with the outcome for any *t*. Coefficient functions for curves aligned using motion were not considered due to the low R-squared.

### 4.1. Windowed Thresholding

The proportional thresholding procedure used to create the sequence of thresholded networks (Section 2.2 and Figure 1), and thus the function-valued network diagnostic curves, has the potential drawback that weaker edges are always analyzed in tandem with stronger edges. This process can conceal organizational patterns that are important to executive function but that occur only in the weakest edges. Therefore, we also considered an alternative *windowed thresholding* approach, which partitions the edges into bins based on their weight (Bassett *and others*, 2012; Lohse *and others*, 2014; Schwarz and McGonigle, 2011). Within each bin, the edges form a network on which the network diagnostic can be calculated, and the network diagnostic curve is a function of the bins (e.g., the bin’s end- or mid-points).

As in Bassett *and others* (2012), we considered windows based on percentile bins, so that a windowed threshold *s* = 0.01 corresponds to the top percentile of edges by strength, *s* = 0.02 is the next highest percentile of edges, and so on. In contrast to proportional-threshold modularity curves, windowed-threshold modularity curves showed that all but the strongest edges had a modularity close to 0 (Figure S.11). The participation coefficient curves in the overall sample appeared similar under both types of thresholding, with age-related patterns similar to those observed in proportional thresholding. Overall, both types of thresholding revealed complementary information: stronger edges exhibited more variation in modularity and participation coefficient between individuals.

## 5. Discussion

In this paper, we represented network models of the brain as functions that describe their multiscale topology. We modeled the association between these function-valued data and executive function, age, and motion through functional regression. To address possible misalignment in the data, we proposed RAFT, a novel approach to combine functional regression with a *supervised* alignment step that incorporates information from auxiliary variables such as age. Through simulations, we investigated the impact of the alignment method on the accuracy of downstream models, thereby demonstrating that supervised alignment often results in functional regression coefficient estimates that are more accurate than—and no worse than—those obtained without any alignment. Moreover, across different simulation settings, RAFT performed well more consistently than the other alignment methods.

The regression coefficient *β*_2_(*t*) describes the relationship between *y* and the network diagnostic curves in their entirety, given the alignment. Namely, *β*_2_(*t*) is the association between variations in *y* and variations in the network curves after removal of phase variation. Among the alignment methods, both RAFT and unsupervised alignment achieved moderate power and coverage for *β*_2_(*t*) given the pointwise confidence intervals of 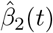 obtained from the functional regression model. No single alignment method consistently out-performed the others in all scenarios, though the *registr* method often had higher power in the presence of sinusoidal noise. Notably, contaminating the covariate curves with sinusoidal noise negatively impacted all methods, even in the ideal case when the true alignment is known. Future analyses incorporating this type of data could involve applying a band-pass filter during data pre-processing. In sensitivity analyses, we discovered that the performance of our proposed methods was sensitive to the warping of the template curve, but was more stable to the choice of template and to uniform warping of all the covariate curves. This pattern of findings suggests that choosing a template which is not representative of the data can severely impact performance; however, overall perturbations of the data appear less likely to affect accuracy. We also found that the good performance of the alignment methods extended to new data with differently shaped covariates (Figure S.3) and that employing simpler covariate functions could achieve similar performance in many cases (Figure S.2). These findings indicate that our overall alignment strategy is flexible and can be successfully adapted to different types of misaligned data.

Although the unsupervised methods sometimes worked well in simulations, they did not improve prediction of executive function in the PNC data. When *z* encoded age as the auxiliary variable, age-aligned RAFT curves on their own were more predictive of executive function compared to the unaligned curves and all other alignment methods (Figure 5). However, we note that this excellent performance may be due to the relatively high correlation between *z* and *y* (*ρ* = 0.64)—any warping of the data designed to maximize prediction accuracy of *z* will naturally also predict *y* well.

Therefore, we investigated if curves that were aligned using *z* to reflect motion as the auxiliary variable would improve prediction of executive function. Unlike age, motion is very weakly correlated with executive function (*ρ*= -0.13), but we anticipated that motion-aligned curves would still improve prediction, as previous studies have established the impact of motion on FC estimates (Power *and others*, 2012). However, this was not the case, and all alignment methods (including no alignment) led to low predictive power in the downstream model (Figure 5), perhaps because motion showed only slight nonlinear and linear associations with the network diagnostics (Figure S.13 and Figure S.14). Therefore, warping the curves did not benefit from any additional information from the motion variable. Another reason could be the limited signal from the FC curves, especially when *t >* 0.5. Curves with flat regions, such as the right tail of both the modularity and participation coefficient curves, could have led to identifiability issues during warping, since in flat regions there are multiple (infinitely many) warping functions that can give rise to the same alignment.

While the main subject of our analysis was network diagnostic curves obtained from a cumulative, proportional thresholding approach, we had also repeated our analyses in curves that were generated by windowed thresholding. The resulting prediction performance was similar to that of the proportional-threshold curves (Figure S.12): namely, the functional data themselves could not explain most of the variability in executive function, unless aligned by age.

There are several limitations of our method. First and most importantly, the model for *z* in Step 2 of the supervised alignment method is inherently overfit. This overfitting derives from the fact that the warping functions are flexible enough that, even with heavy penalization, a small perturbation in the functional predictors can result in predictions 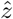 that perfectly overlap with the observed *z*. It is for this reason that we do not use the outcome of interest *y* itself to warp the curves, as explained in Section 2.4. Consequently, curves aligned using *z* should not be used in downstream analyses where *z* is the outcome of interest. This observation also means that the variable *z* should be chosen such that its correlation with *y* is substantially less than 1. (In simulations, we found that correlations of up to around 0.8 still resulted in acceptable estimation of the true association *β*_2_(*t*) between *y* and the aligned curves.) If *y* and *z* are similar, then aligned curves can predict *y* well, even without a true association (Figure 5). This prediction is possible because the curves are aligned such that they best predict *z*, and if *y* and *z* are similar, then they will also predict *y*. Thus, our method is most useful when *z* is not associated with *y*, but is associated with the warping process. In that case, we can be more confident that any significant estimate of *β*_2_(*t*) in Model (2.4) indicates a true association between the curves and *y*.

Second, the functions 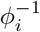 have at most one inflection point and *ξ*(*t*) have none; these may be too simple to align data with multiple features and complex misalignment mechanisms. In that case, B-splines or spline-like methods that join multiple CDF functions together at pre-determined knots may be a useful extension. Third, interpretation of the regression coefficients requires care. Due to overfitting, we do not recommend interpreting the *β*_1_(*t*) coefficient functions used to predict the auxiliary variable *z*. However, the coefficient function *β*_2_(*t*) can still provide meaningful insights, provided that interpretations are conditioned on the alignment (i.e., the choice of template, and the auxiliary variable used to align them.) Finally, we proposed alignment as a possible solution to the problems outlined in (van den Heuvel *and others*, 2017); we did not compare our method to other pre-processing models that address the reasons for systematic differences in the distribution of functional connectivity edge weights (Afyouni *and others*, 2019). Our work has several straightforward extensions and applications to other domains. While phase variation was considered a nuisance in the analysis of network curves, it can be a quantity of interest in other applications, particularly in time series analysis. For instance, in diurnal activity data obtained from accelerometers, the delayed onset of wake or sleep can manifest in phase variability in the activity curves, and is often an important outcome related to chronotype, age, and disease. While previous studies have considered scalar-on-function regression (Leroux *and others*, 2019) and phase variability (McDonnell *and others*, 2021) separately in the context of activity measurement, our method can account for these together. For instance, age can be used as an auxiliary variable to align wake-sleep cycles across individuals in order to better predict disease status. At the same time, the degree of misalignment can be related to circadian dysfunction and overall health. As the warping functions 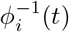 are themselves functional data, and may be related to the outcome of interest, we could consider including them in an additional term in Model (2.2):

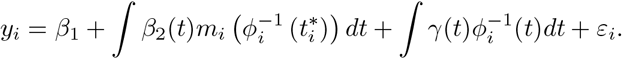

In that case, the coefficient function *γ*(*t*) would represent the association between the warping function for person *i* and their outcome, given the aligned curves. Another mode of future work could involve the estimation of a single warping function for two or more functional predictors simultaneously, such as *m*_*i*_(*t*) and *p*_*i*_(*t*), assuming that the warping mechanism is the same across network diagnostics. As alignment is highly relevant to time-varying data with periodic trends, future work could involve the application of our method to alternate domains, such as accelerometry and activity measurements from wearables (McDonnell *and others*, 2021). Finally, the analysis of the derivatives of the modularity and participation coefficient curves could give us further insight into brain network development in two ways: first, the rate of change over thresholds may be an important feature related to executive function. Second, integrating and differentiating the curves may provide additional landmarks that can help with alignment. As yet, the intersection of topological data analysis and functional data analysis is still not fully explored, and we look forward to more work at this fascinating juncture.

## 6. Software

The R code implementation of the proposed methods, both the unsupervised Beta CDF warping and RAFT, is publicly available on Github: https://github.com/danni-tu/RAFT.

## 7. Supplementary Material

Supplementary material is available online at http://biostatistics.oxfordjournals.org.

## Acknowledgments

The authors thank Haochang Shou for helpful discussions and Ellyn Butler for help with data acquisition. This work was supported by the NIH grants R01 MH112847, R37 MH125829, R01 MH113550, and R01 MH120482. The Philadelphia Neurodevelopmental Cohort was supported by the National Institute of Mental Health grants MH089983 and MH089924.

---

**Fig. S.1.**
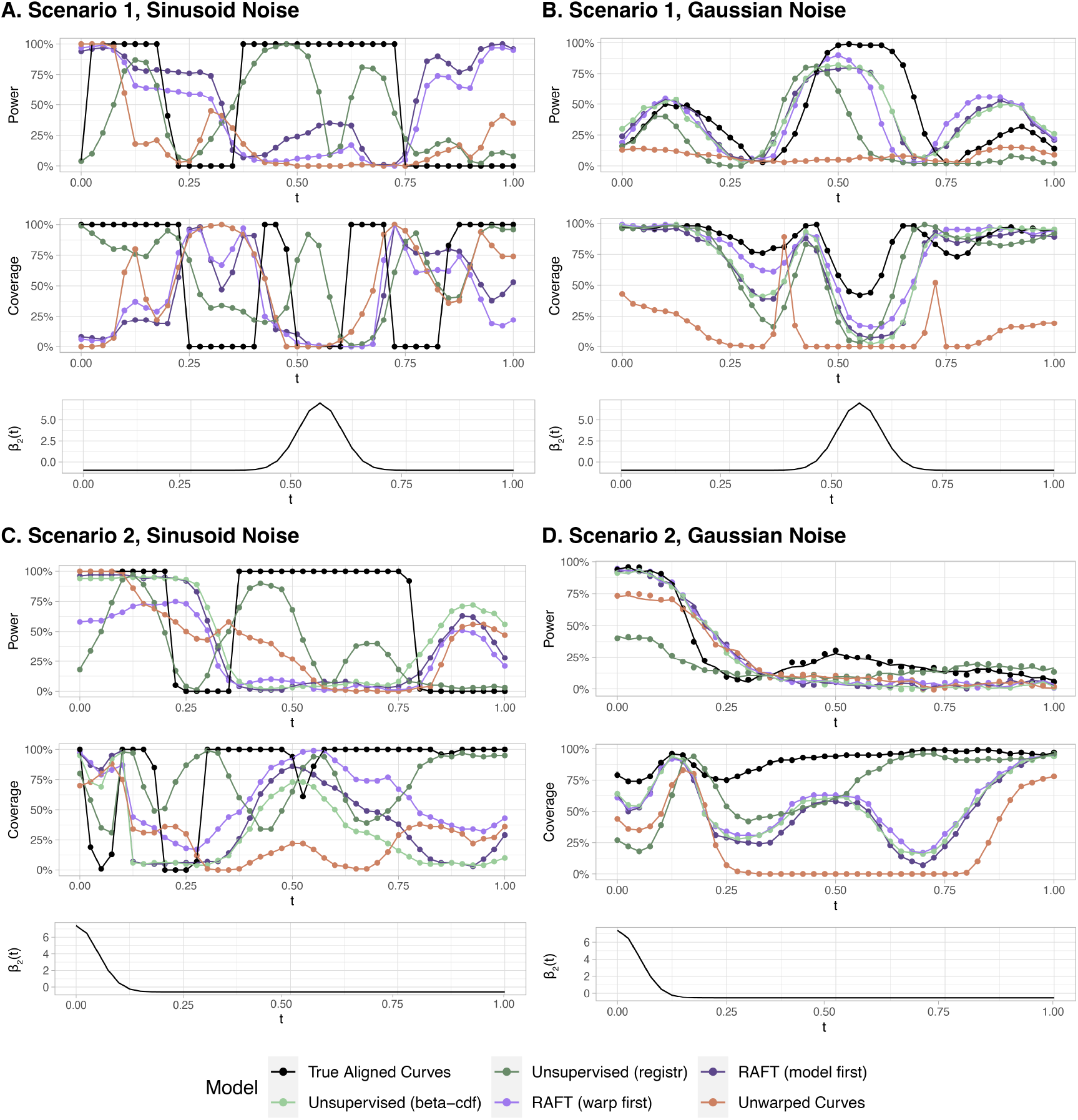
Alignment methods achieved moderate power and coverage using the pointwise confidence interval for 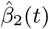 obtained from the penalized functional regression model fit on the aligned curves. For each *t*, power was defined as the proportion of repetitions where the confidence interval excluded 0 if *β*_2_(*t*)*≠* 0, and coverage as the proportion of repetitions where the confidence interval included the true value of *β*_2_(*t*). In most cases, any alignment improved power and coverage beyond that of no alignment, with *registr* generally performing best in curves with added sinusoidal noise, and our proposed methods performing best in curves with added Gaussian noise. Note: in Panel D, many methods had similar performance and a 2% vertical jitter was added to the data points in order to distinguish between them.

**Fig. S.2.**
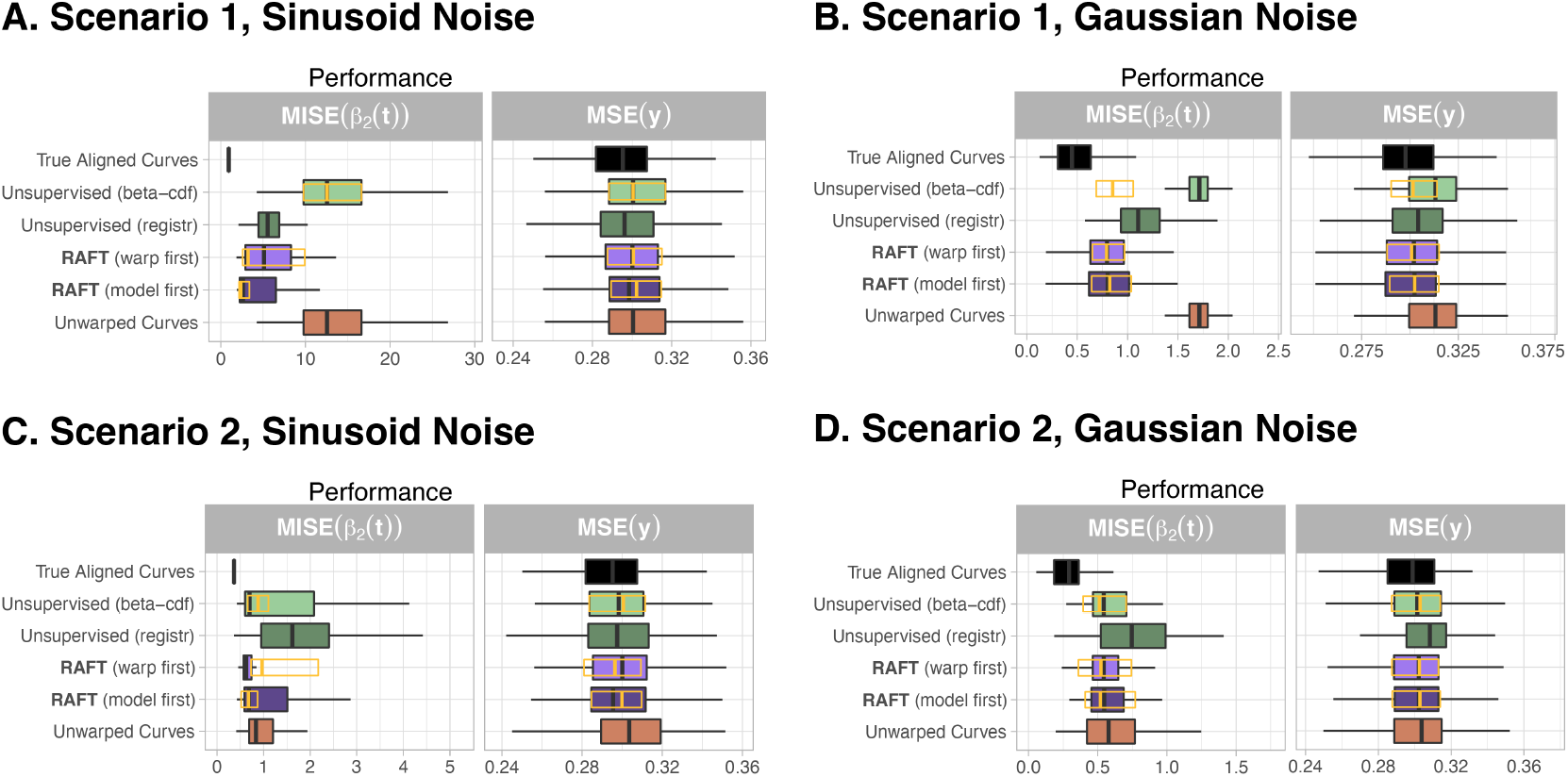
In simulated data, alignment methods employing simpler polynomial warping functions (Section 2.5) can also improve the accuracy of model coefficient estimation beyond using the unwarped curves. Aside from the warping functions, the data-generating scenarios in Panels A to D are identical to those described in Section 3.1. For comparison, the orange outline represents the 25th, 50th, and 75th percentile of performance achieved in our main results using Beta CDF warping functions in Figure 3.

**Table S1.**
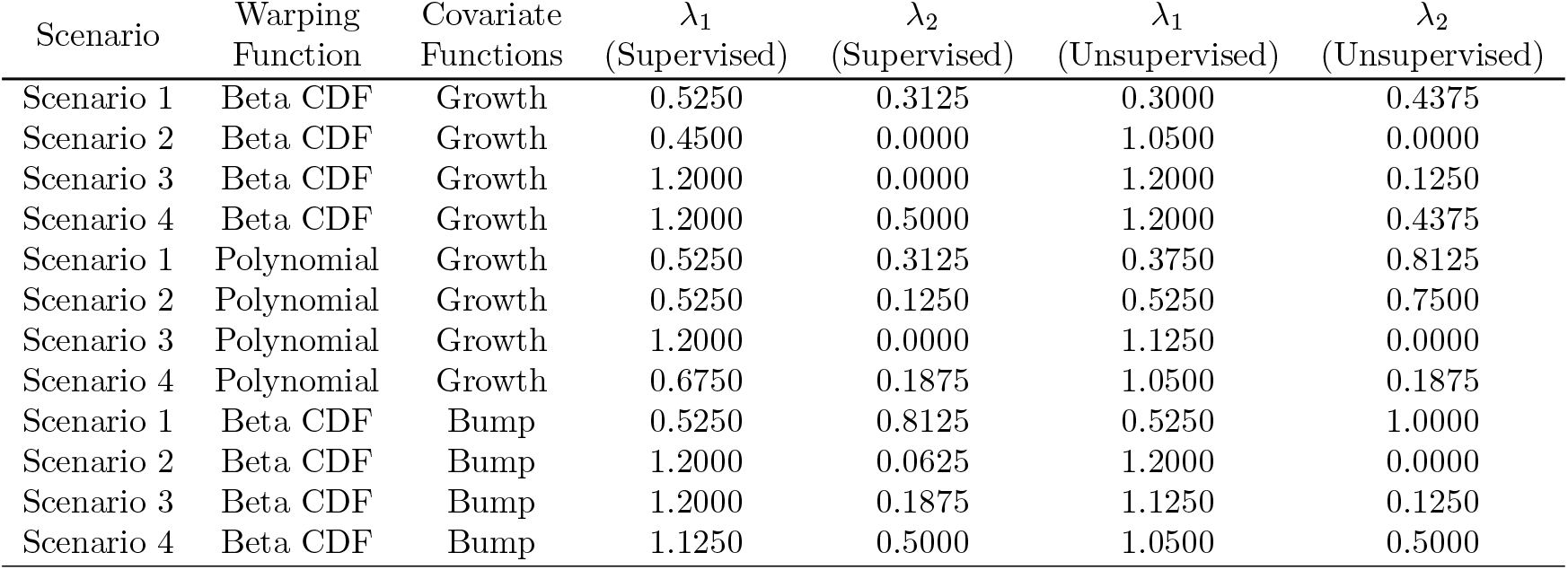
Penalty parameters used in estimating the aligned functions using the loss function in (2.3). These were calculated separately for each data generating scenario described in Section 3.1 and sensitivity analyses, as well as for the supervised and unsupervised algorithms. Penalty parameters were estimated using a grid search (Section 2.6.1) on a data set of *n* = 125 generated curves that were separate from the testing data used to evaluate performance.

**Fig. S.3.**
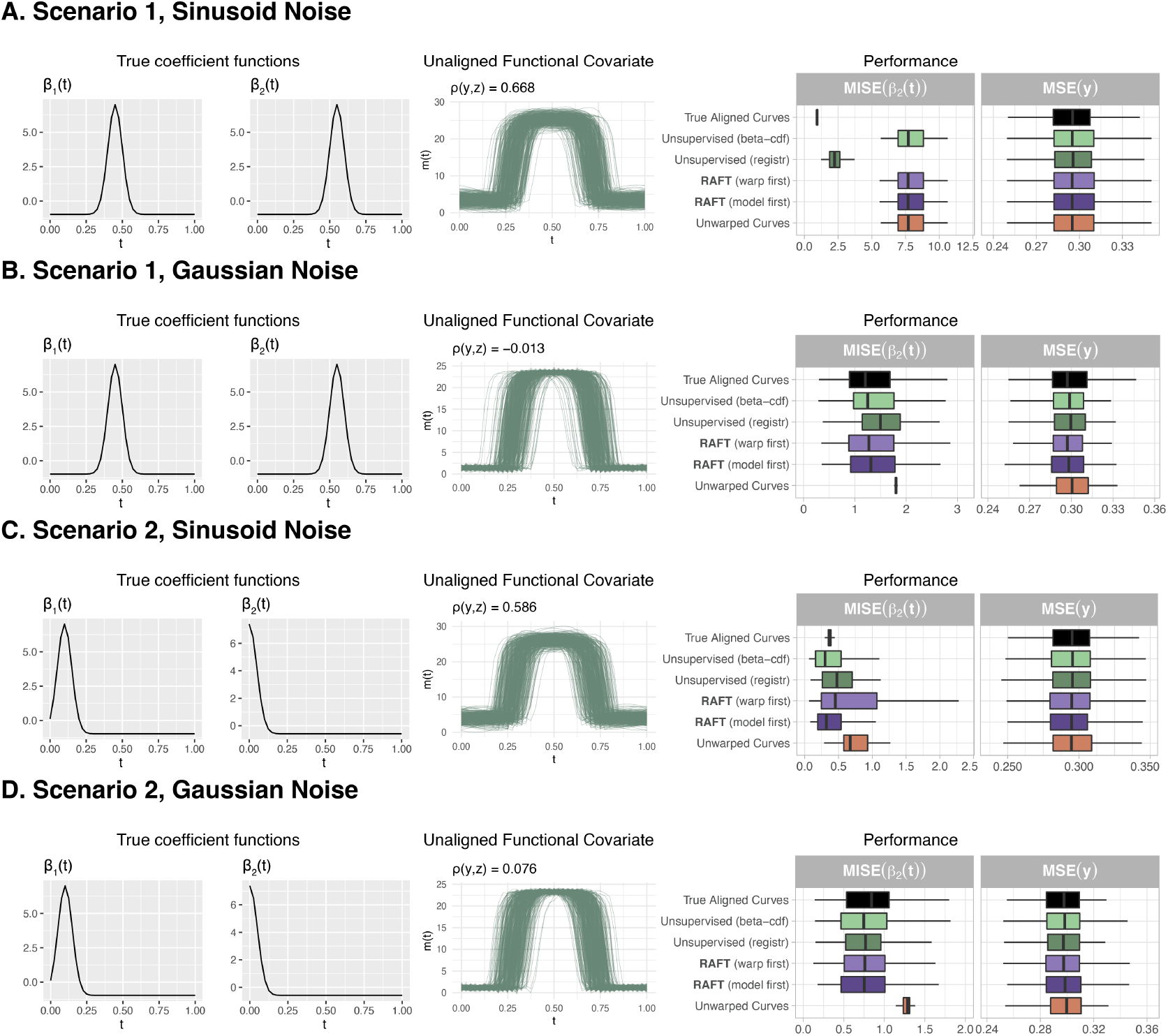
In another set of simulated data with bump-shaped covariate functions mimicking diurnal activity profiles, alignment can improve the accuracy of model coefficient estimation beyond using the unwarped curves. For these types of covariate functions, *registr* performed similarly to or better than the unsupervised and RAFT algorithms. Interestingly, for the scenario described in Panel A involving the first set of true *β*_1_(*t*) and *β*_2_(*t*) functions and covariate curves with added sinusoidal noise, the estimated value of *λ*_2_ was high enough that no warping was done in either the supervised or the unsupervised cases, resulting in identical performance for 4 of the 6 methods.

**Fig. S.4.**
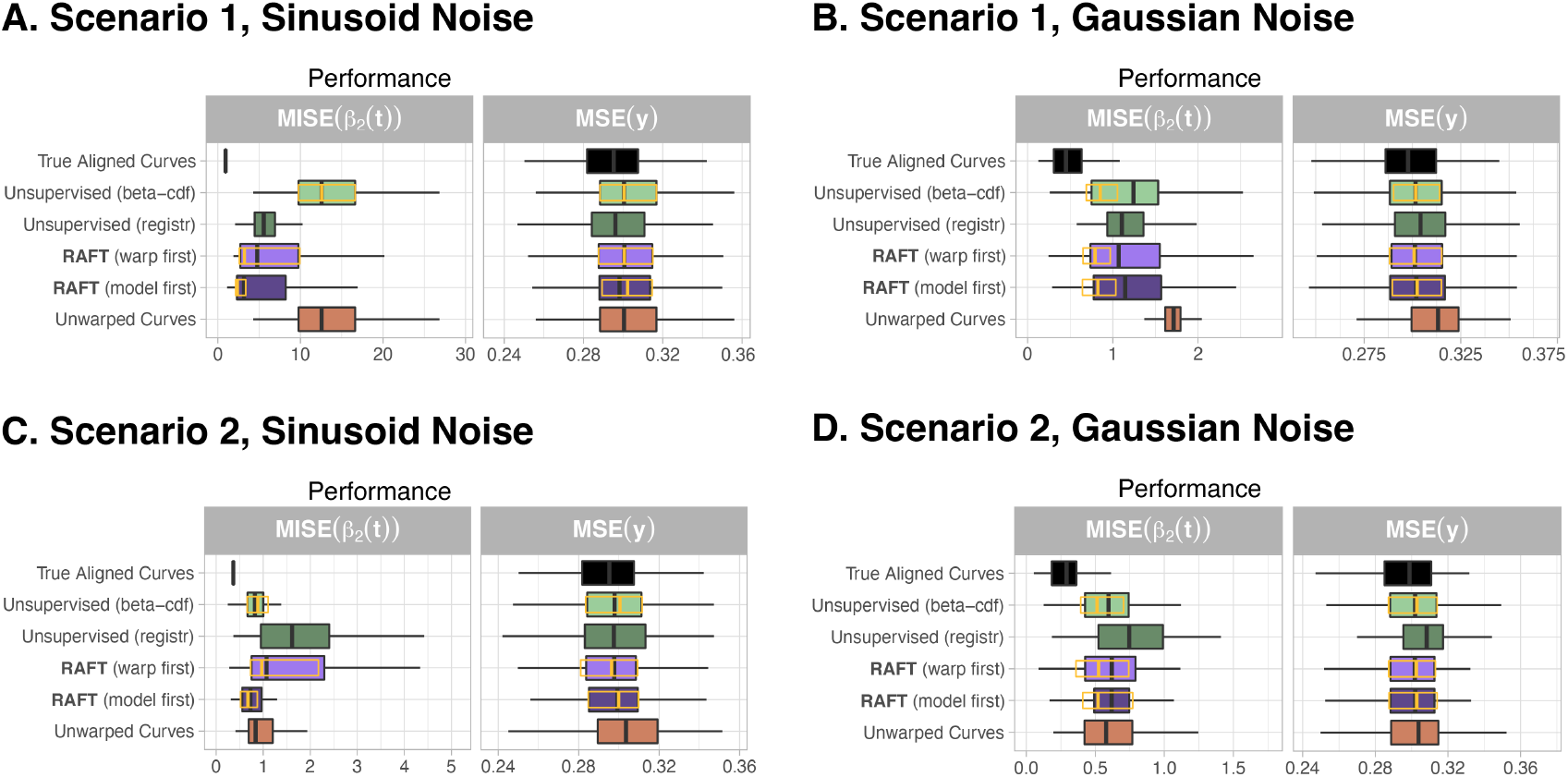
Performance remains comparable or is slightly poorer when the template function is randomly chosen from the observations, as opposed to the default *L*^2^ centroid. For comparison, the orange outline represents the 25th, 50th, and 75th percentile of performance achieved in our main results in Figure 3.

**Fig. S.5.**
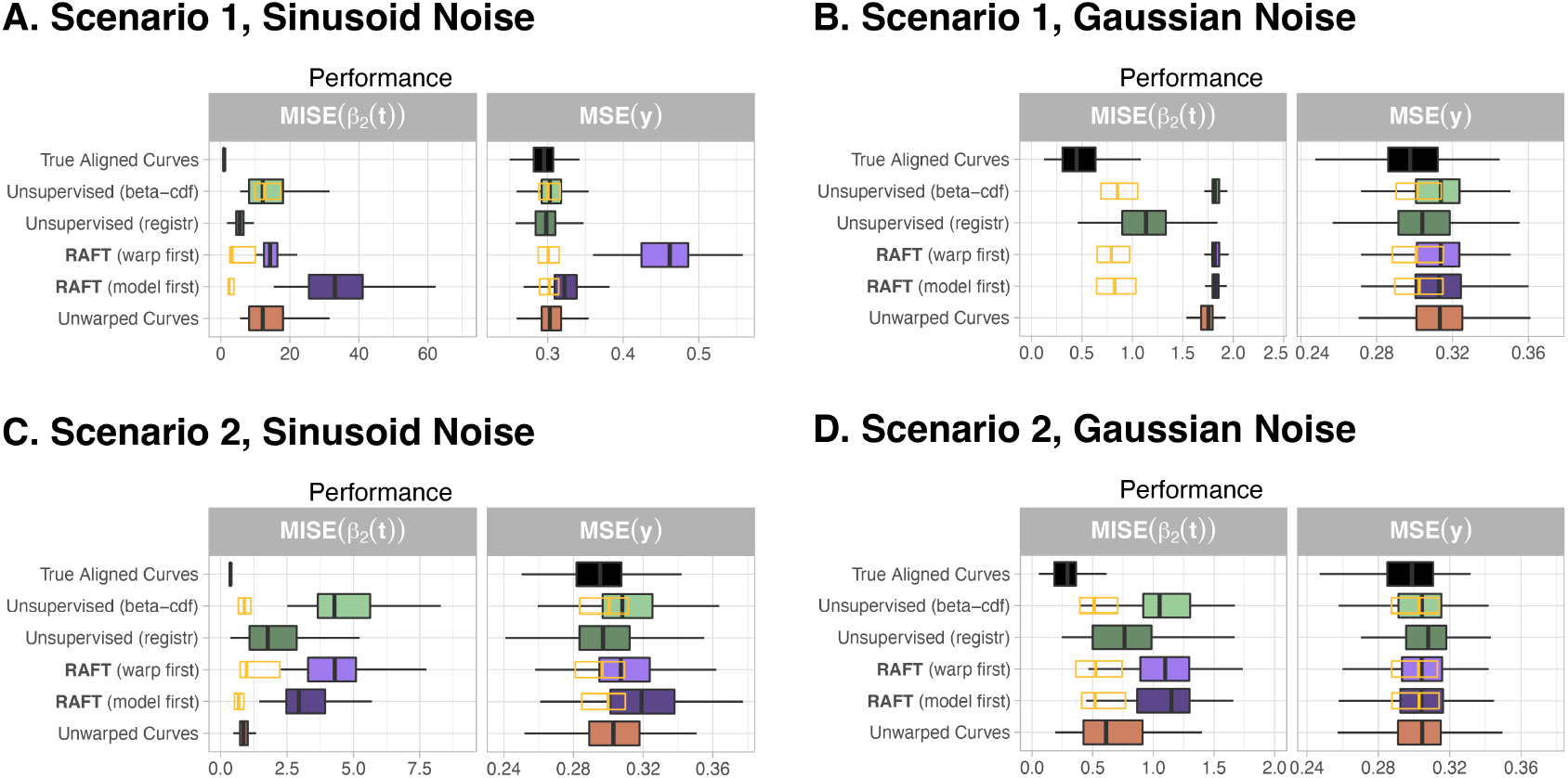
When the *L*^2^ centroid is used as the template, but is further warped by a known quantity so that it becomes less representative of the other data, the performance of algorithms relying on that template degrades. In most scenarios, the estimation error of RAFT and the unsupervised methods have now exceeded that of using no alignment. For comparison, the orange outline represents the 25th, 50th, and 75th percentile of performance achieved in our main results in Figure 3.

**Fig. S.6.**
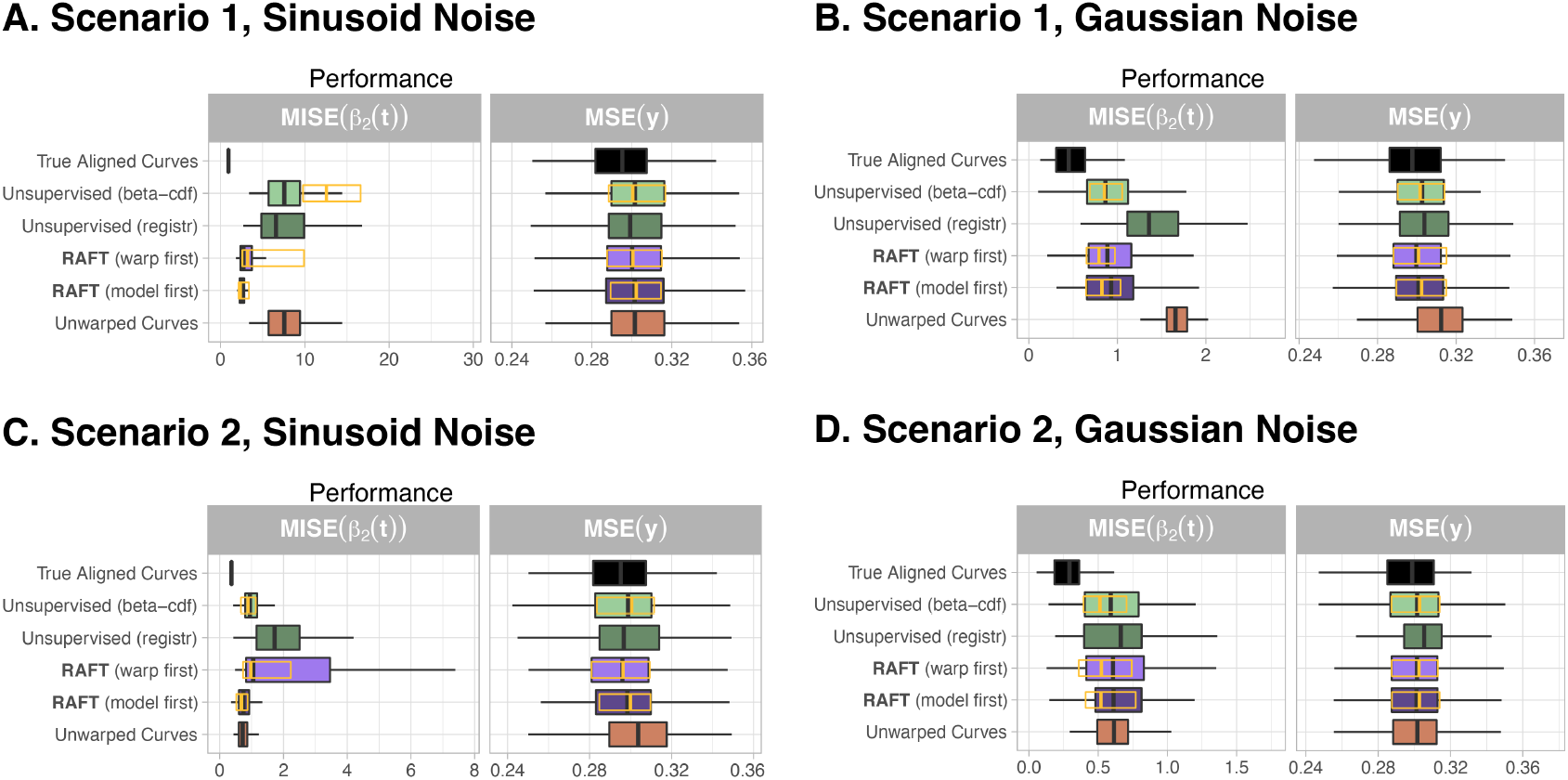
When all of the covariate data have been uniformly warped, both the supervised and unsupervised methods perform as well as, if not better than, the same methods on the original simulation data. For comparison, the orange outline represents the 25th, 50th, and 75th percentile of performance achieved in our main results in Figure 3.

## S.1. Philadelphia Neurodevelopmental Cohort Data

### S.1.1 Participants

The Philadelphia Neurodevelopmental Cohort is a publicly available multi-platform study of child development between the ages of 8 and 21 years (Satterthwaite *and others*, 2016). A collaborative NIMH-funded effort between the Children’s Hospital of Philadelphia and the Brain Behavior Laboratory at the University of Pennsylvania, the PNC study incorporates structural neuroimaging, resting-state and task-based functional neuroimaging, and psychiatric and neurocognitive assessments. All adults and guardians of minors provided informed consent, and the study procedures were approved by the Institutional Review Board of the University of Pennsylvania and the Children’s Hospital of Philadelphia.

#### S.1.2. Neurocognitive Battery

The Penn Computerized Neurocognitive Battery (CNB) comprises 14 separate tests designed to assess accuracy and speed across a range of cognitive domains including executive control, episodic memory, and social cognition (Gur *and others*, 2012). Factor loadings of general cognitive efficiency (which combines speed and accuracy across all constructs) were obtained from a confirmatory bifactor analysis and used as a score of cognitive performance (Moore *and others*, 2015) after linear scaling by the mean and standard deviation.

#### S.1.3. Functional Connectivity Data

Resting-state functional MRI (rs-fMRI) data was collected from *n* = 1601 participants from a single 3T Siemens TIM Trio whole-body scanner with a 32-channel head coil using a whole-brain, single-shot, multi-slice, gradient-echo echoplanar sequence with a voxel resolution of 3 *×* 3 *×* 3 mm. Sequence parameters were as follows: time repetition = 3000 ms, time echo = 32 ms, field of view = 192 *×* 192 mm, matrix = 64 *×* 64 for 46 slices, slice thickness = 3 mm, slice gap = 0 mm, and flip angle = 90 degrees. Further details can be found in Satterthwaite *and others* (2014)

Pre-processing of the BOLD time series included a confound regression procedure to reduce noise and motion artifacts, and was implemented using the eXtensible Connectivity Pipeline (XCP) Engine (Ciric *and others*, 2017). The BOLD time series were then mapped to the cortical surface (Tooley *and others*, 2019; Cui *and others*, 2020) and averaged over 400 areas of the cortex as described in Schaefer *and others* (2017). The 400-area parcellation was selected as it preserves the structure of the 7 and 17 communities from Yeo *and others* (2011) while compromising between spatial resolution and dimensionality. We considered only the cortical regions as they demonstrate significant reorganization and integration throughout development (Fair *and others*, 2009; Baum *and others*, 2017; Tooley *and others*, 2019), although subcortical and cerebellar regions are important to understanding brain changes in adolescence (Gu *and others*, 2015). Of the *n* = 1601 participants who underwent multimodal neuroimaging, those with missing resting data (*n* = 196), more than 20 frames with motion exceeding 0.25 mm (*n* = 45), or a mean relative RMS displacement higher than 0.2 mm per frame (*n* = 7) were excluded (Satterthwaite *and others*, 2013), resulting in a final imaging sample of *n* = 1123 participants. Of those, *n* = 1122 had complete neurocognitive and imaging data.

**Fig. S.7.**
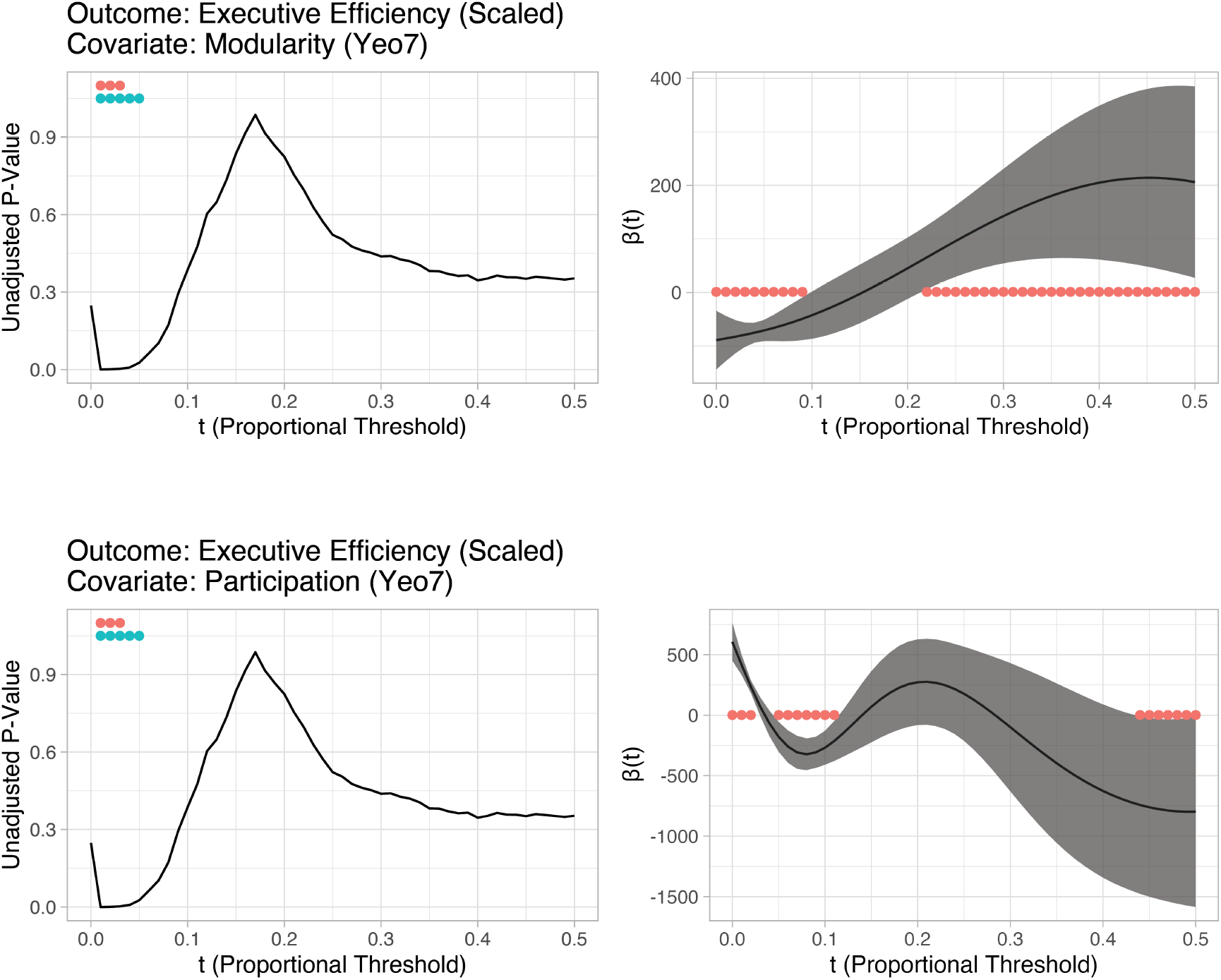
Functional regression methods can capture the relationship between network diagnostics, age, and executive function jointly over values of threshold *t*, thereby resulting in greater statistical power to detect significant associations between network diagnostics and executive function compared to multiple pointwise analyses, even if the latter do not perform multiplicity correction. Left column: Unadjusted *p*-values for the linear model with covariates *m*_*i*_(*t*) and age, for each value of *t*. Right column: The estimated coefficient *β*(*t*) of the scalar-on-function regression model with functional covariate *m*_*i*_(*t*) (unaligned) and age as a scalar covariate. The red dots correspond to regions of *m*_*i*_(*t*) significantly associated with the outcome (i.e., thresholds *t* where the 95% confidence interval for *β*(*t*) does not contain 0).

**Fig. S.8.**
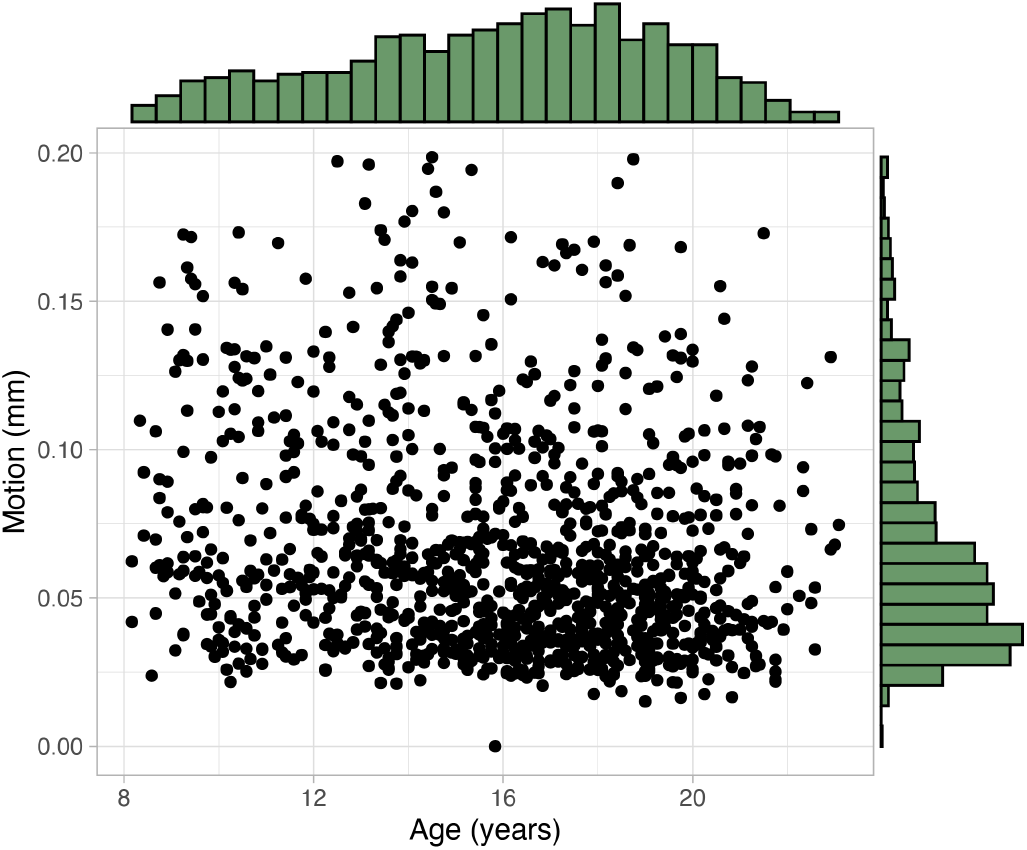
Scatterplot and marginal histograms of age and motion variables from the Philadelphia Neurodevelopmental Cohort. There was a weak negative relationship between age and motion (*ρ*=0.167, 95% CI: [-0.223,-0.110]).

**Table S2.**
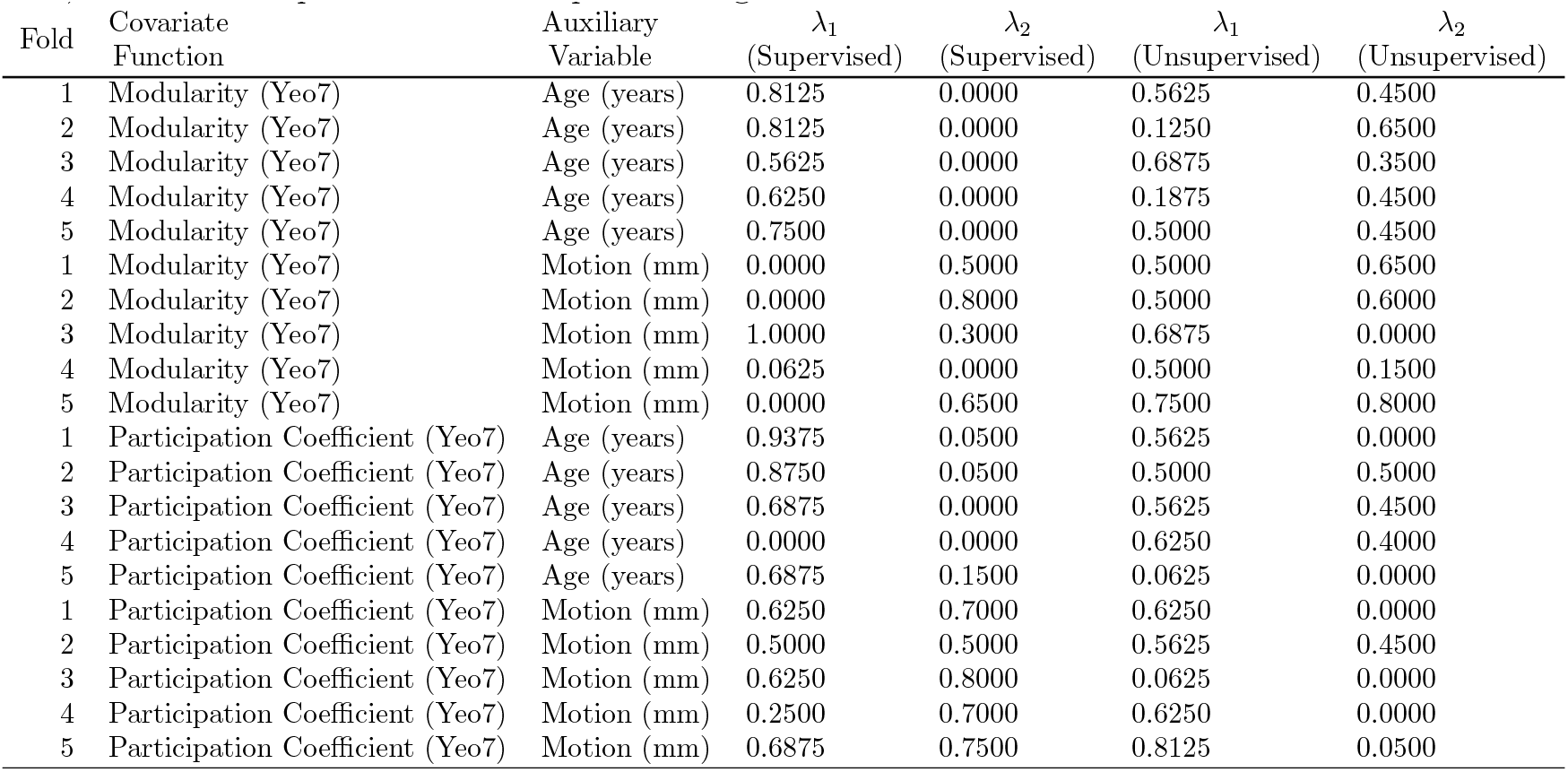
Penalty parameters used in estimating the aligned functions in the PNC data using the loss function in (2.3). In order to assess performance on all of the observations in the PNC data, the data were partitioned into 5 equally-sized folds. The parameters were estimated using a grid search (Section 2.6.1) for each fold and performance was evaluated on the remaining folds, similar to 5-fold cross-validation. Therefore, the values of *λ*_1_ and *λ*_2_ were calculated separately for each fold, covariate function, auxiliary variable, and for the supervised and unsupervised algorithms.

**Fig. S.9.**
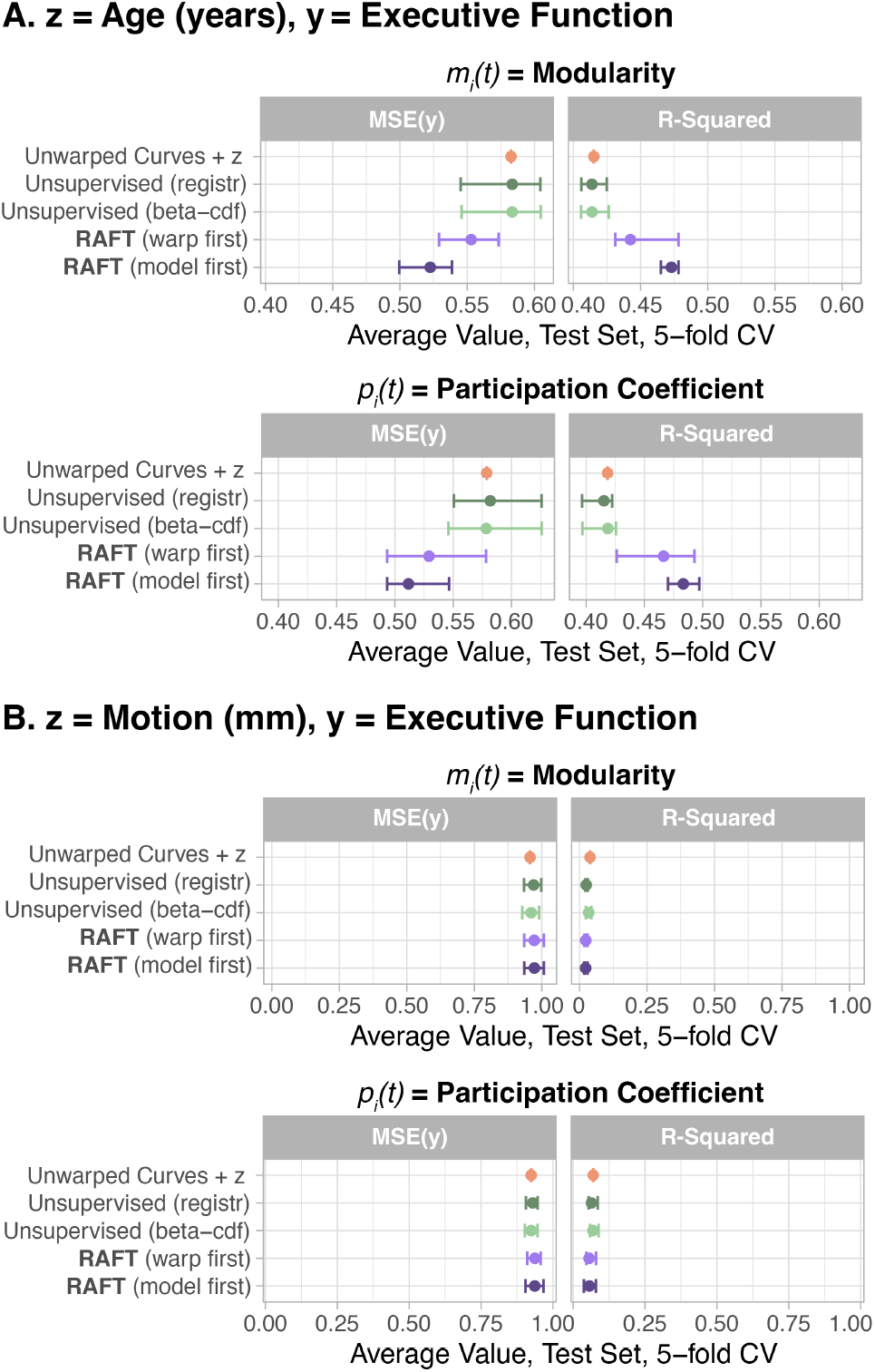
Comparison of alignment methods and unaligned curves in scalar-on-function regression models with a linear term for when *z* reflected age (Panel A) or motion (Panel B). When *z* reflected age, RAFT performs well against other methods. But when *z* reflected motion, all methods perform similarly, indicating that alignment of the network diagnostic curves adds little to the predictive power beyond that given by the linear term.

**Fig. S.10.**
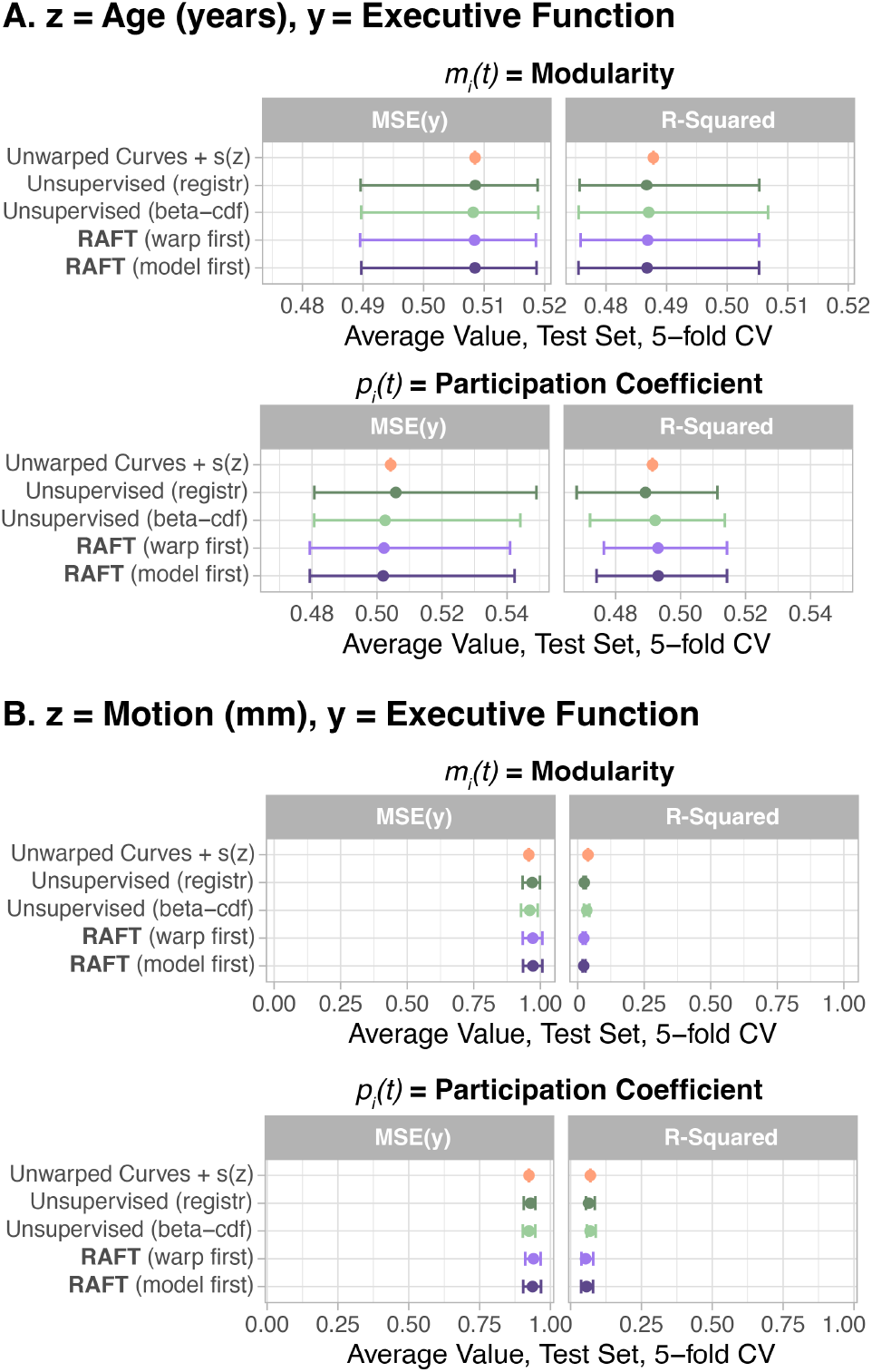
Comparison of alignment methods and unaligned curves in scalar-on-function regression models with a nonlinear term *s*(*z*) for when *z* reflected age (Panel A) or motion (Panel B). When *z* reflected age, the average performance of all methods is similar, indicating that age-supervised alignment of the network diagnostic curves adds little to the predictive power beyond that given by the *s*(*z*) term.

**Fig. S.11.**
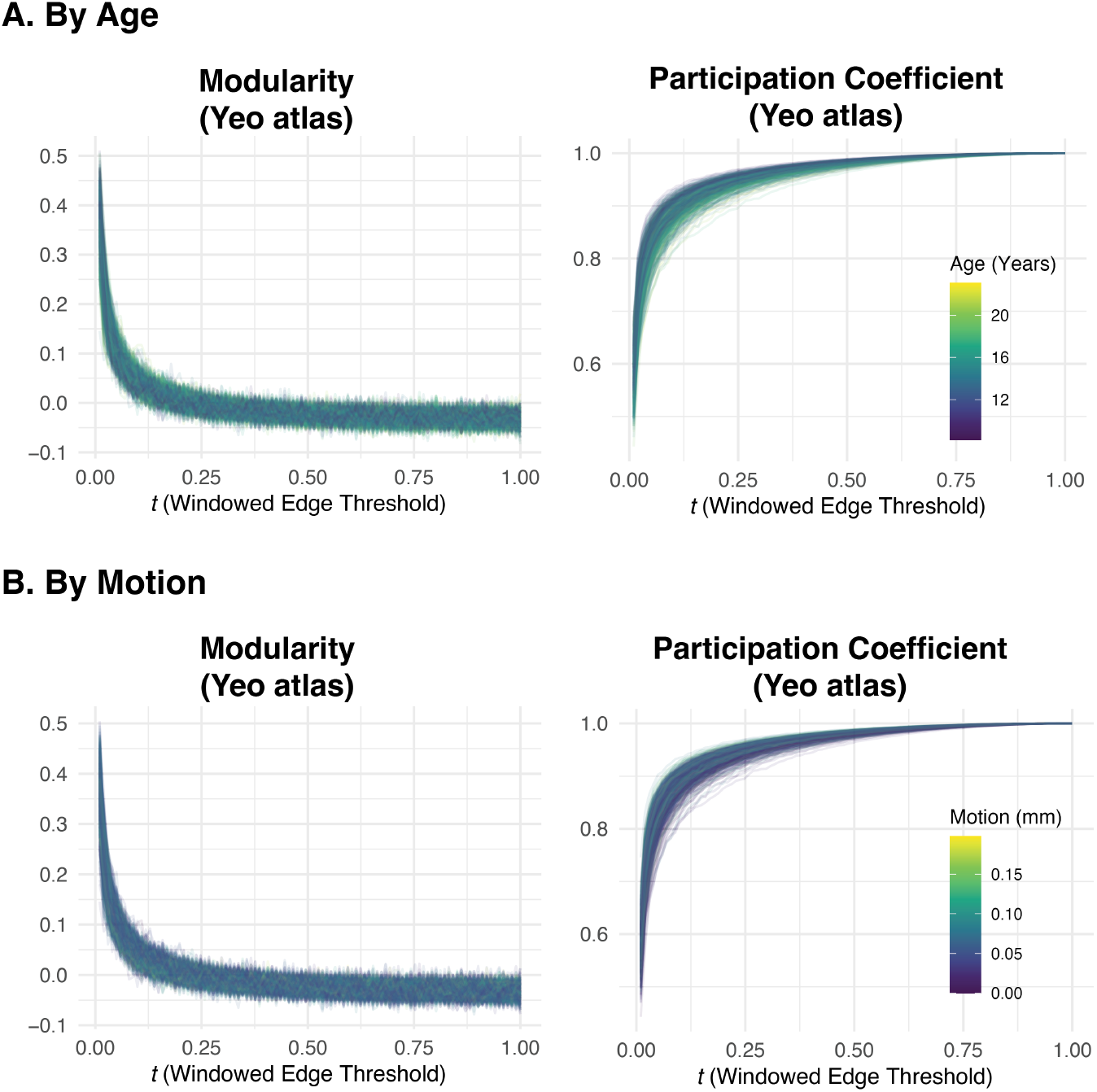
Data from the Philadelphia Neurodevelopmental Cohort (Satterthwaite *and others*, 2016) using the alternate windowed thresholding approach. Modularity and participation coefficient were calculated given the 7 communities defined in Yeo *and others* (2011) as a function of the percentile *t*, where *t* = 0.01 corresponds to the strongest percentile of edges, *t* = 0.02 corresponds to the next percentile, and so forth. Compared to those obtained from proportional thresholding, window-threshold modularity curves were noisier for *t >*= 0.2 (i.e., the weakest 80 percentile bins), likely because weaker edges have a more random configuration. In contrast, the participation coefficient curves appeared more similar for both thresholding types. The color of the curve corresponds to age (Panel A; darker = younger) or motion (Panel B; darker = less motion).

**Fig. S.12.**
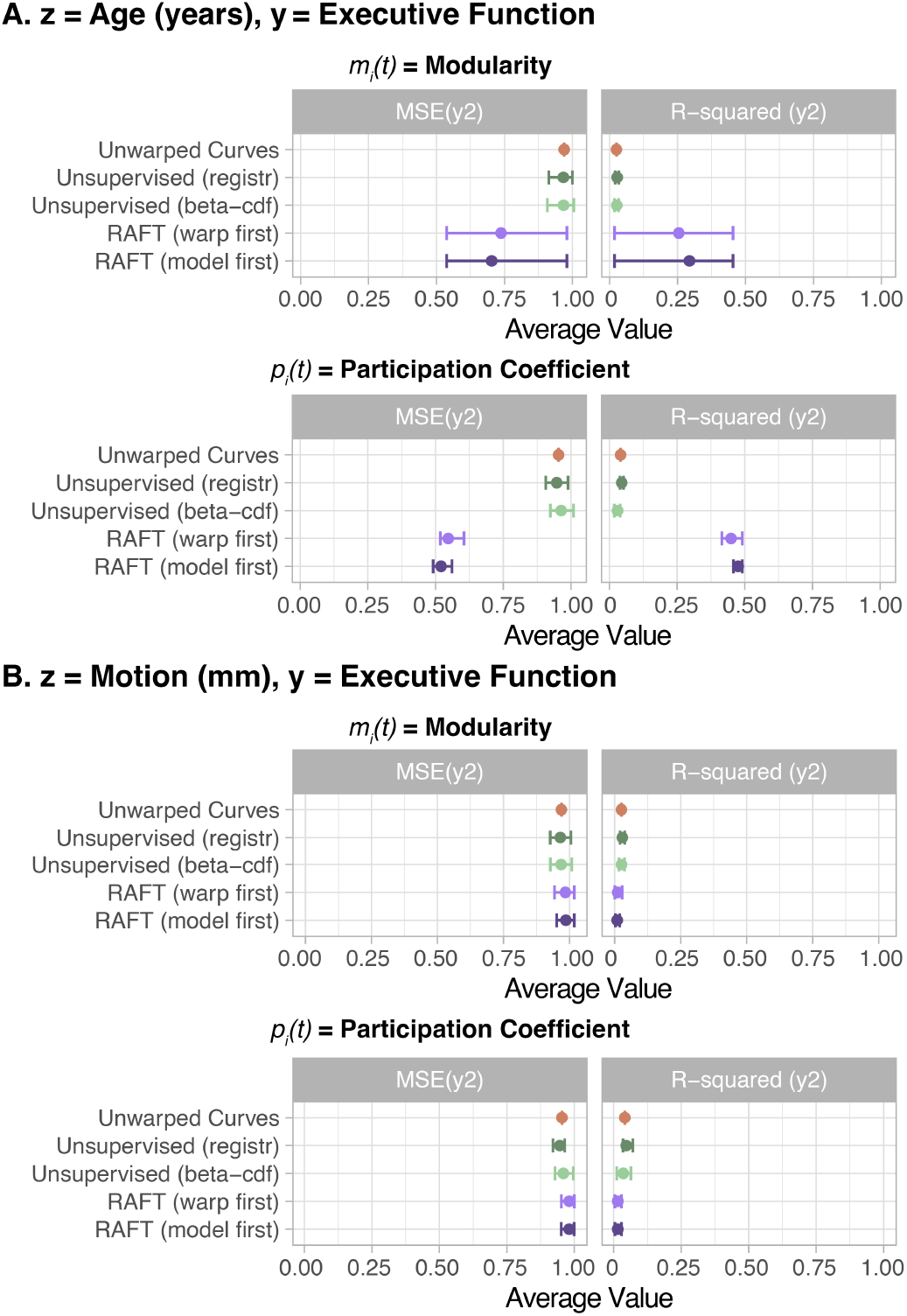
In data from the Philadelphia Neurodevelopmental Cohort (Satterthwaite *and others*, 2016), thresholded using the alternate windowed approach, RAFT alignment of the modularity and participation coefficient curves improved the prediction of executive function when the auxiliary variable was age, but not when the auxiliary variable was motion. We considered modularity (left column) and participation coefficient (right column) using communities defined by the 7-community parcellation described in (Yeo *and others*, 2011); age was measured in years, and motion was measured as average relative RMS displacement in millimeters. As with the proportional-threshold curves, RAFT performed better when *z* reflected age (Panel A) compared to when *z* reflected motion (Panel B), likely for similar reasons.

**Fig. S.13.**
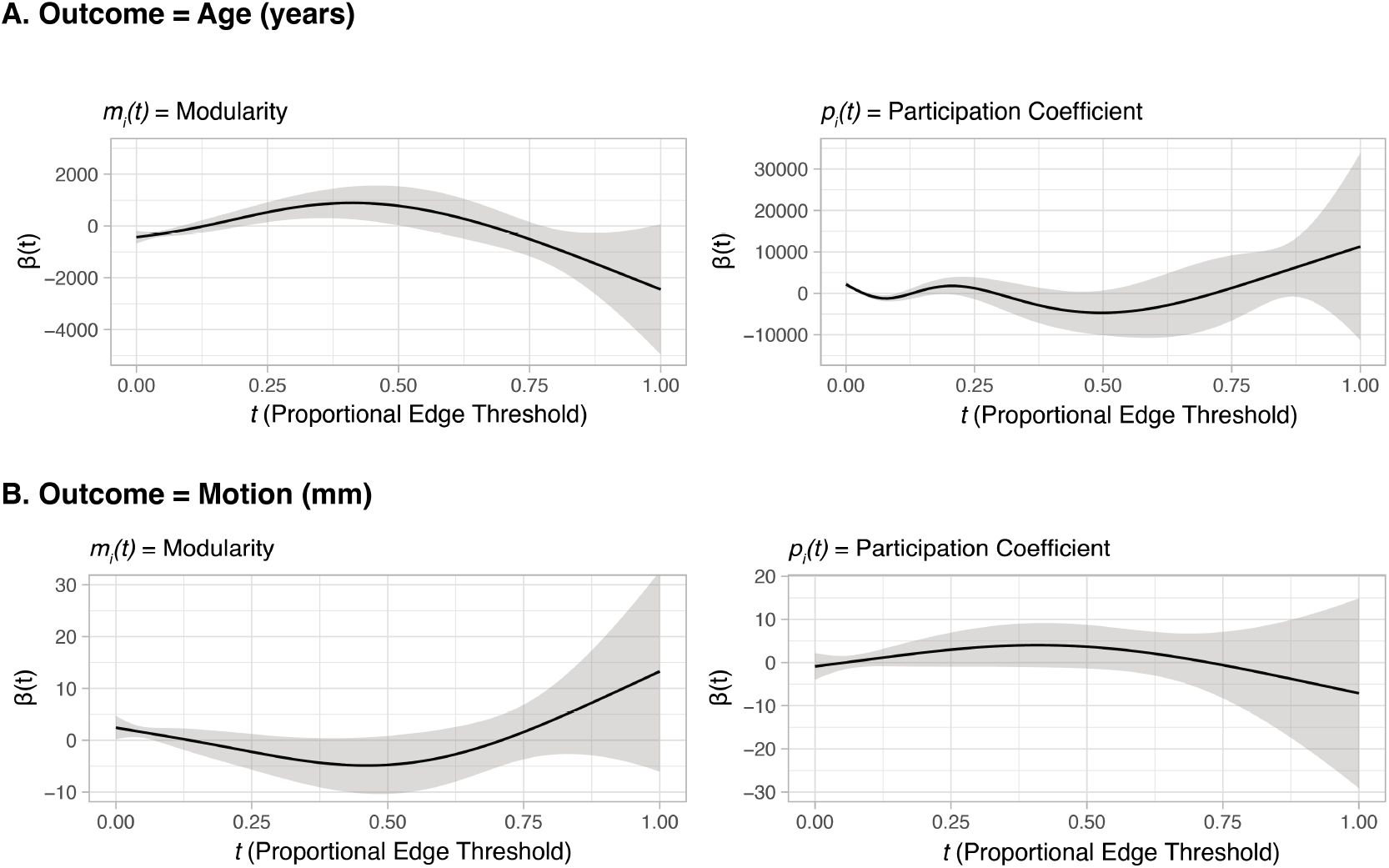
Non-linear, conditional associations between the auxiliary variables *z* and the *unwarped* network diagnostic curves in their entirety. (Panel A) Scalar-on-function regression coefficients where age is the outcome and the predictor is either modularity (left) or participation coefficient (right). We found significant associations between age and modularity curves for *t <* 0.5, and between age and participation coefficient curves for *t <* 0.3. (Panel B) Scalar-on-function regression coefficients where motion is the outcome and the predictor is either modularity (left) or participation coefficient (right). We found significant associations between motion and modularity for *t <* 0.15, and we did not observe any significant associations between motion and participation coefficient.

**Fig. S.14.**
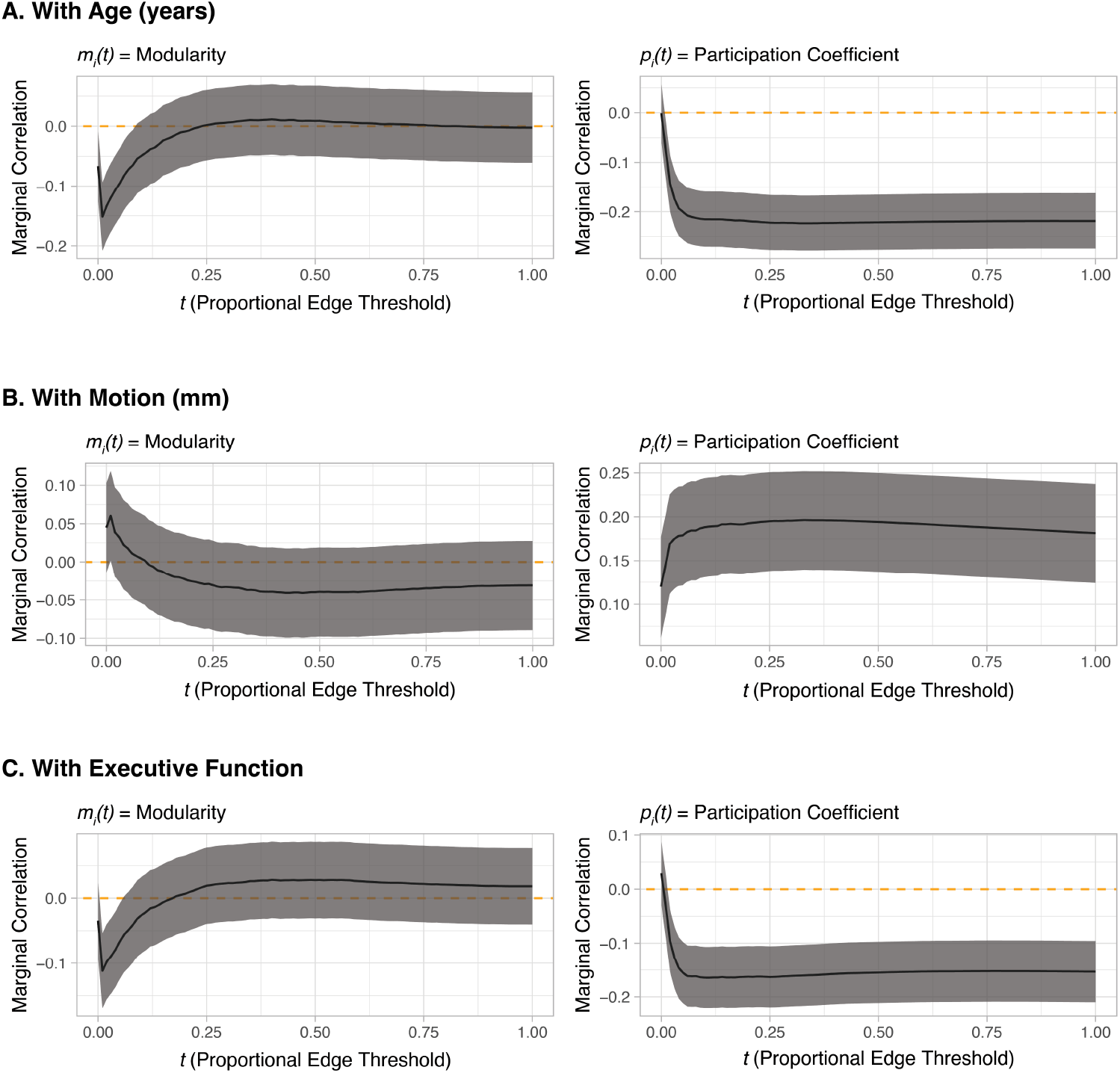
Marginal linear relationships (correlations) between the auxiliary variables *z* or executive function *y*, and the network diagnostic curves at each threshold *t*. The correlations and their 95% percent confidence intervals were calculated separately for each value of *t*. (Panel A) The marginal correlation, as a function of *t*, between age and modularity (left), and between age and participation coefficient (right). (Panel B) The marginal correlation, as a function of *t*, between motion and modularity (left), and between motion and participation coefficient (right). (Panel C) The marginal correlation, as a function of *t*, between executive function and modularity (left), and between executive function and participation coefficient (right). This figure and Figure S.13 contain complementary information: While the *non-linear* associations in Figure S.13 are between the scalar variables and the entire curves (i.e., jointly over all *t*), these are *linear* associations between the scalar variables and the network diagnostics at a fixed value of *t*, independent of all other values of *t*.

**Fig. S.15.**
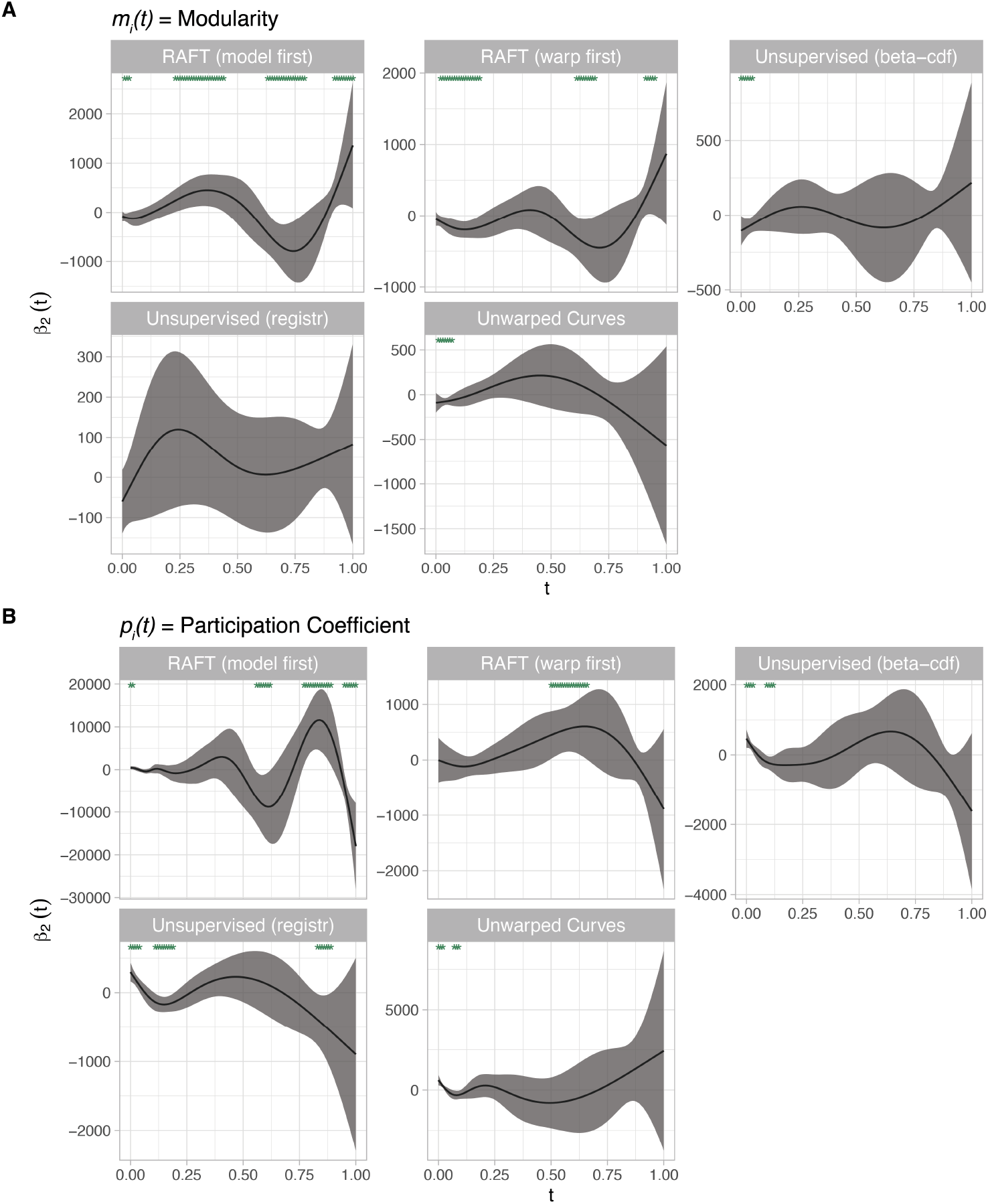
Estimated 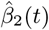 from Model 2.2 representing the association between executive function and modularity (Panel A) or participation coefficient (Panel B) after alignment by *z* = age. The unsupervised and supervised models used the penalty parameters obtained from the cross-validation step (fold 3) provided in Table S.2. Asterisks at the top of the panel signal regions of the functional domain *t* where 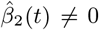. Note that these coefficient functions each correspond to a different set of aligned covariate curves.

