## Supplement for "Regression and Alignment for Functional Data and Network Topology"

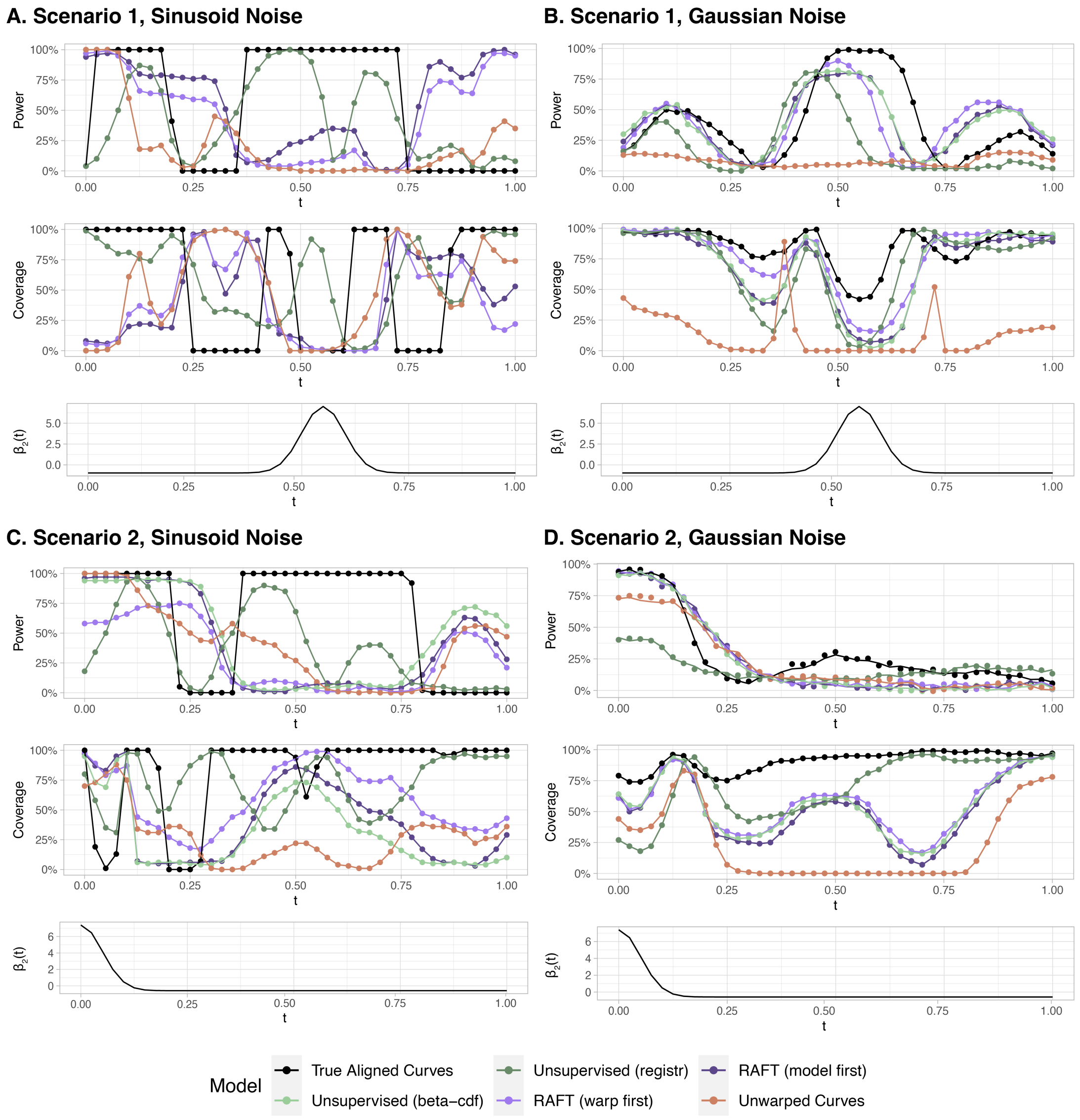


Fig. S.1. Alignment methods achieved moderate power and coverage using the pointwise confidence interval for $\hat{\beta}_2(t)$ obtained from the penalized functional regression model fit on the aligned curves. For each $t$, power was defined as the proportion of repetitions where the confidence interval excluded 0 if $\beta_2(t) \neq 0$, and coverage as the proportion of repetitions where the confidence interval included the true value of $\beta_2(t)$. In most cases, any alignment improved power and coverage beyond that of no alignment, with *registr* generally performing best in curves with added sinusoidal noise, and our proposed methods performing best in curves with added Gaussian noise. Note: in Panel D, many methods had similar performance and a 2% vertical jitter was added to the data points in order to distinguish between them.

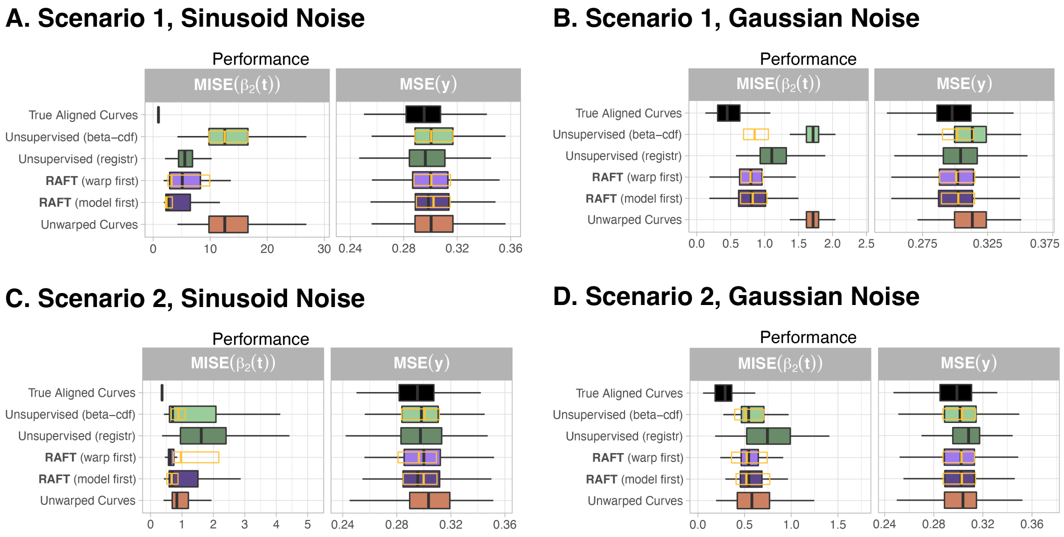


Fig. S.2. In simulated data, alignment methods employing simpler polynomial warping functions (Section 2.5) can also improve the accuracy of model coefficient estimation beyond using the unwarped curves. Aside from the warping functions, the data-generating scenarios in Panels A to D are identical to those described in Section 3.1. For comparison, the orange outline represents the 25th, 50th, and 75th percentile of performance achieved in our main results using Beta CDF warping functions in Figure 3.

Table S.1. Penalty parameters used in estimating the aligned functions using the loss function in (2.3). These were calculated separately for each data generating scenario described in Section 3.1 and sensitivity analyses, as well as for the supervised and unsupervised algorithms. Penalty parameters were estimated using a grid search (Section 2.6.1) on a data set of $n = 125$ generated curves that were separate from the testing data used to evaluate performance.

| Scenario | Warping Function | Covariate Functions | $\lambda_1$ (Supervised) | $\lambda_2$ (Supervised) | $\lambda_1$ (Unsupervised) | $\lambda_2$ (Unsupervised) |
|---|---|---|---|---|---|---|
| Scenario 1 | Beta CDF | Growth | 0.5250 | 0.3125 | 0.3000 | 0.4375 |
| Scenario 2 | Beta CDF | Growth | 0.4500 | 0.0000 | 1.0500 | 0.0000 |
| Scenario 3 | Beta CDF | Growth | 1.2000 | 0.0000 | 1.2000 | 0.1250 |
| Scenario 4 | Beta CDF | Growth | 1.2000 | 0.5000 | 1.2000 | 0.4375 |
| Scenario 1 | Polynomial | Growth | 0.5250 | 0.3125 | 0.3750 | 0.8125 |
| Scenario 2 | Polynomial | Growth | 0.5250 | 0.1250 | 0.5250 | 0.7500 |
| Scenario 3 | Polynomial | Growth | 1.2000 | 0.0000 | 1.1250 | 0.0000 |
| Scenario 4 | Polynomial | Growth | 0.6750 | 0.1875 | 1.0500 | 0.1875 |
| Scenario 1 | Beta CDF | Bump | 0.5250 | 0.8125 | 0.5250 | 1.0000 |
| Scenario 2 | Beta CDF | Bump | 1.2000 | 0.0625 | 1.2000 | 0.0000 |
| Scenario 3 | Beta CDF | Bump | 1.2000 | 0.1875 | 1.1250 | 0.1250 |
| Scenario 4 | Beta CDF | Bump | 1.1250 | 0.5000 | 1.0500 | 0.5000 |

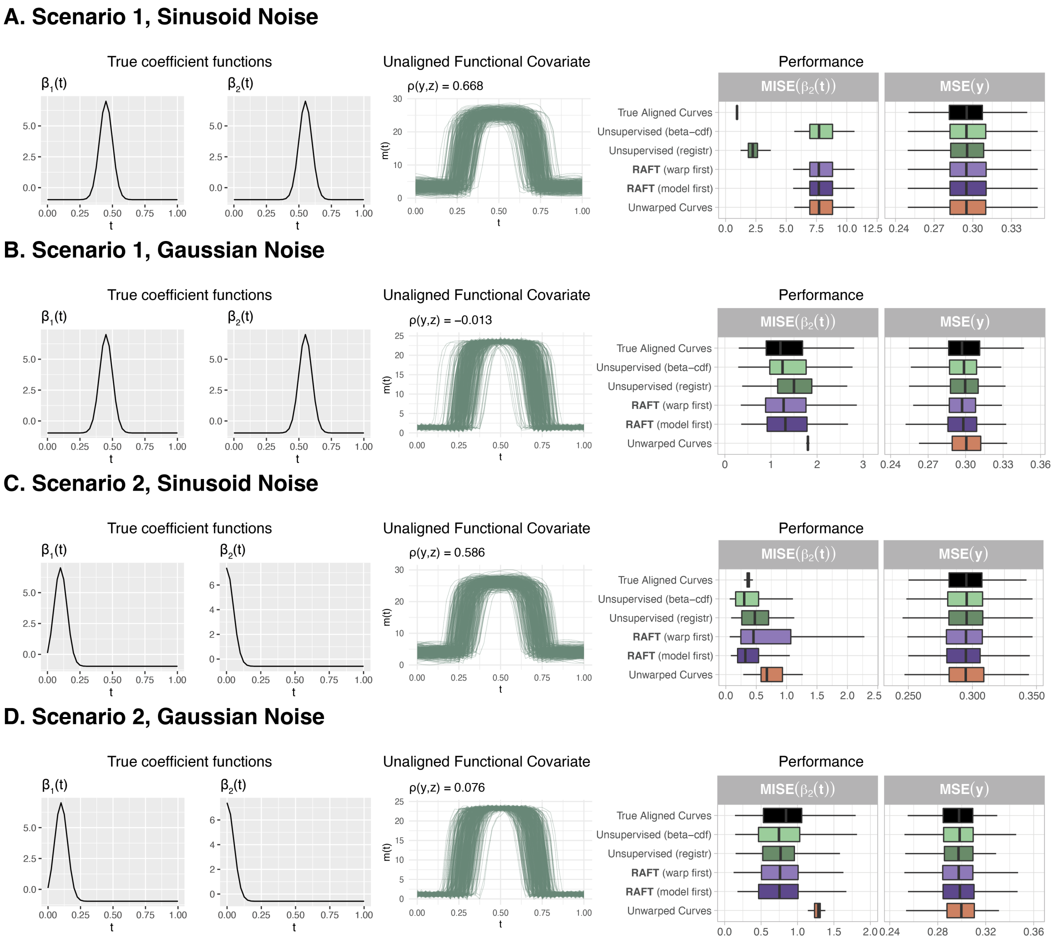


Fig. S.3. In another set of simulated data with bump-shaped covariate functions mimicking diurnal activity profiles, alignment can improve the accuracy of model coefficient estimation beyond using the unwarped curves. For these types of covariate functions, *registr* performed similarly to or better than the unsupervised and RAFT algorithms. Interestingly, for the scenario described in Panel A involving the first set of true $\beta_1(t)$ and $\beta_2(t)$ functions and covariate curves with added sinusoidal noise, the estimated value of $\lambda_2$ was high enough that no warping was done in either the supervised or the unsupervised cases, resulting in identical performance for 4 of the 6 methods.

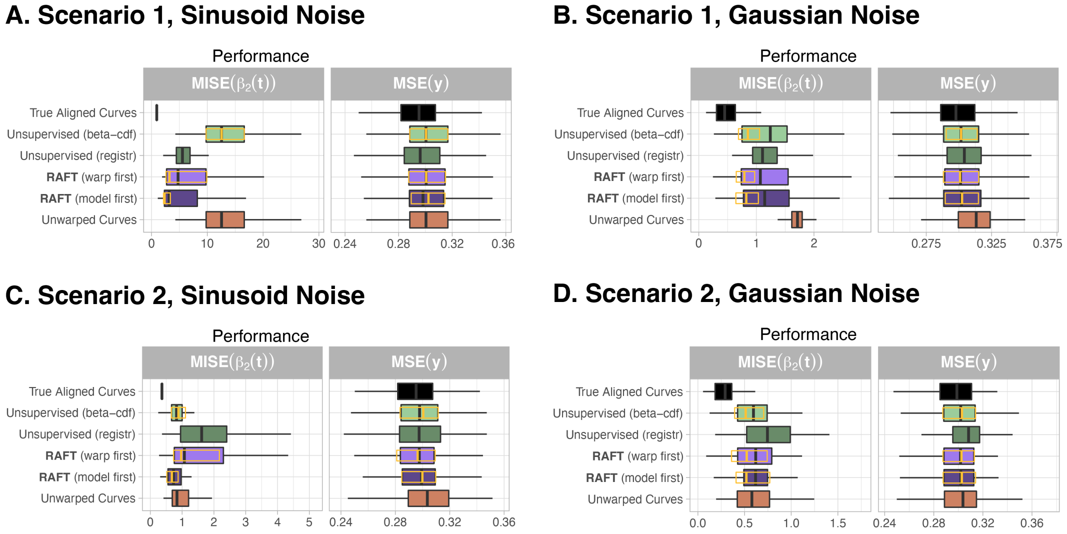


Fig. S.4. Performance remains comparable or is slightly poorer when the template function is randomly chosen from the observations, as opposed to the default $L^2$ centroid. For comparison, the orange outline represents the 25th, 50th, and 75th percentile of performance achieved in our main results in Figure 3.

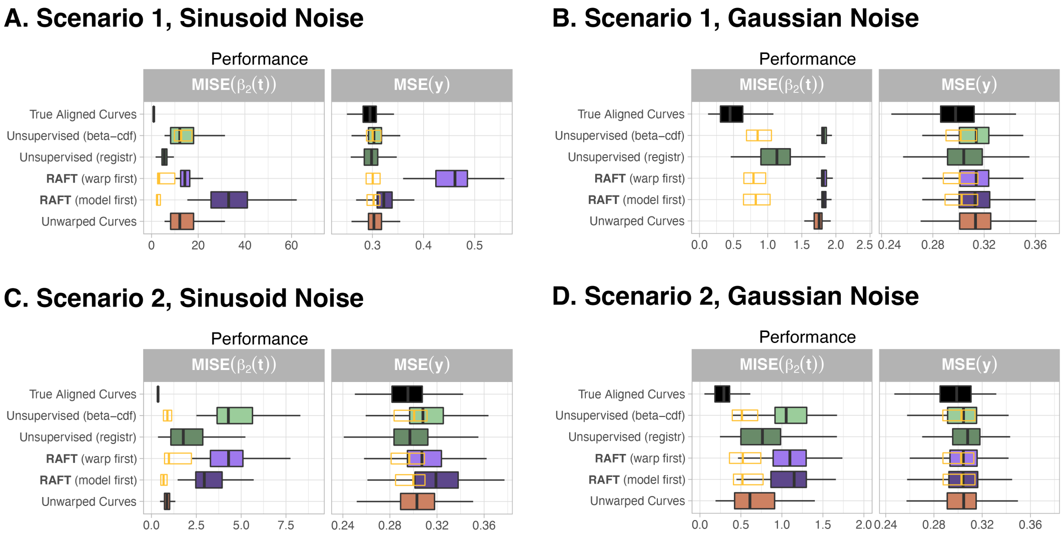


Fig. S.5. When the $L^2$ centroid is used as the template, but is further warped by a known quantity so that it becomes less representative of the other data, the performance of algorithms relying on that template degrades. In most scenarios, the estimation error of RAFT and the unsupervised methods have now exceeded that of using no alignment. For comparison, the orange outline represents the 25th, 50th, and 75th percentile of performance achieved in our main results in Figure 3.

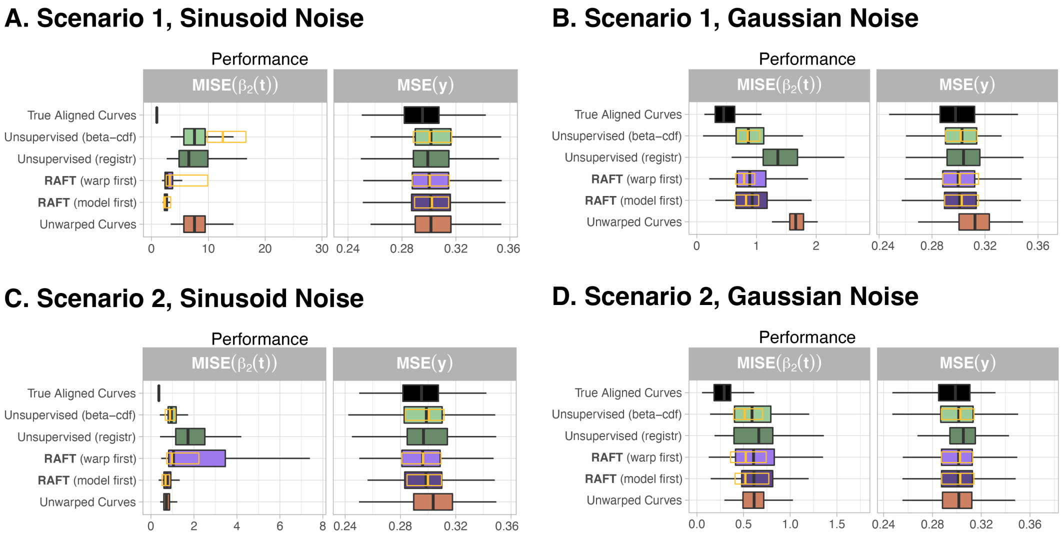


Fig. S.6. When all of the covariate data have been uniformly warped, both the supervised and unsupervised methods perform as well as, if not better than, the same methods on the original simulation data. For comparison, the orange outline represents the 25th, 50th, and 75th percentile of performance achieved in our main results in Figure 3.

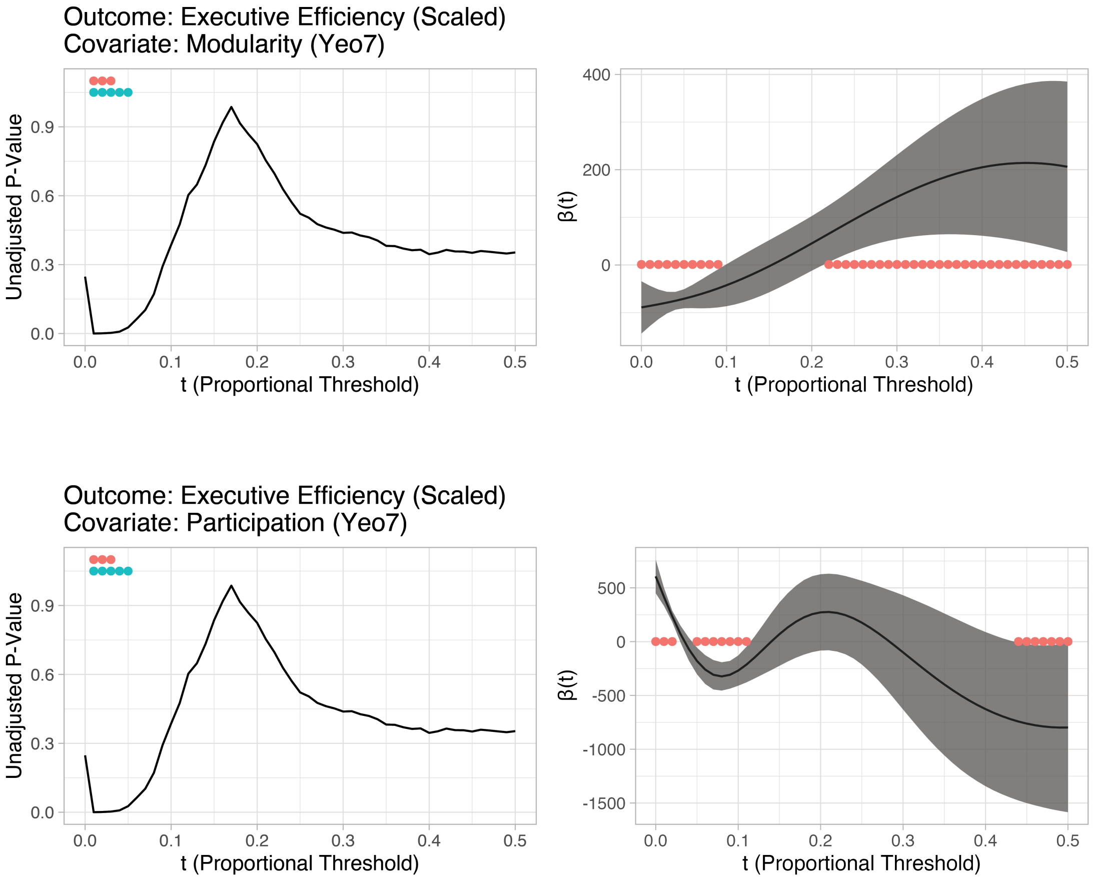


Fig. S.7. Functional regression methods can capture the relationship between network diagnostics, age, and executive function jointly over values of threshold $t$, thereby resulting in greater statistical power to detect significant associations between network diagnostics and executive function compared to multiple pointwise analyses, even if the latter do not perform multiplicity correction. Left column: Unadjusted $p$-values for the linear model with covariates $m_i(t)$ and age, for each value of $t$. Right column: The estimated coefficient $\beta(t)$ of the scalar-on-function regression model with functional covariate $m_i(t)$ (unaligned) and age as a scalar covariate. The red dots correspond to regions of $m_i(t)$ significantly associated with the outcome (i.e., thresholds $t$ where the 95% confidence interval for $\beta(t)$ does not contain 0).

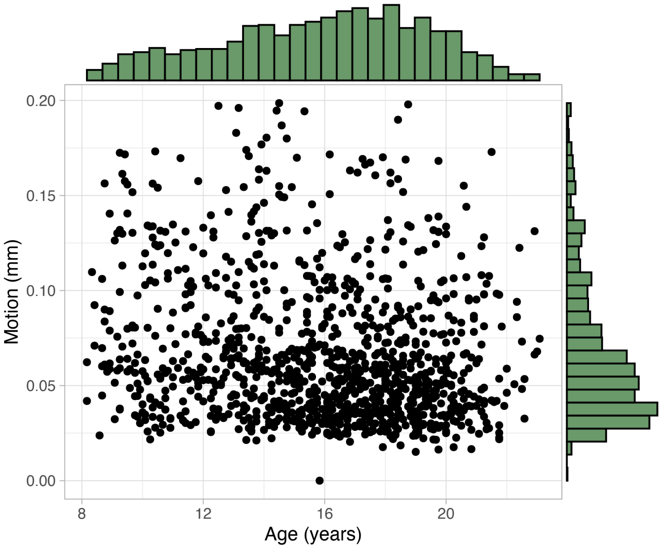


Fig. S.8. Scatterplot and marginal histograms of age and motion variables from the Philadelphia Neurodevelopmental Cohort. There was a weak negative relationship between age and motion ($\rho$=0.167, 95% CI: [-0.223,-0.110]).

Table S.2. Penalty parameters used in estimating the aligned functions in the PNC data using the loss function in (2.3). In order to assess performance on all of the observations in the PNC data, the data were partitioned into 5 equally-sized folds. The parameters were estimated using a grid search (Section 2.6.1) for each fold and performance was evaluated on the remaining folds, similar to 5-fold cross-validation. Therefore, the values of $\lambda_1$ and $\lambda_2$ were calculated separately for each fold, covariate function, auxiliary variable, and for the supervised and unsupervised algorithms.

| Fold | Covariate Function | Auxiliary Variable | $\lambda_1$ (Supervised) | $\lambda_2$ (Supervised) | $\lambda_1$ (Unsupervised) | $\lambda_2$ (Unsupervised) |
|---|---|---|---|---|---|---|
| 1 | Modularity (Yeo7) | Age (years) | 0.8125 | 0.0000 | 0.5625 | 0.4500 |
| 2 | Modularity (Yeo7) | Age (years) | 0.8125 | 0.0000 | 0.1250 | 0.6500 |
| 3 | Modularity (Yeo7) | Age (years) | 0.5625 | 0.0000 | 0.6875 | 0.3500 |
| 4 | Modularity (Yeo7) | Age (years) | 0.6250 | 0.0000 | 0.1875 | 0.4500 |
| 5 | Modularity (Yeo7) | Age (years) | 0.7500 | 0.0000 | 0.5000 | 0.4500 |
| 1 | Modularity (Yeo7) | Motion (mm) | 0.0000 | 0.5000 | 0.5000 | 0.6500 |
| 2 | Modularity (Yeo7) | Motion (mm) | 0.0000 | 0.8000 | 0.5000 | 0.6000 |
| 3 | Modularity (Yeo7) | Motion (mm) | 1.0000 | 0.3000 | 0.6875 | 0.0000 |
| 4 | Modularity (Yeo7) | Motion (mm) | 0.0625 | 0.0000 | 0.5000 | 0.1500 |
| 5 | Modularity (Yeo7) | Motion (mm) | 0.0000 | 0.6500 | 0.7500 | 0.8000 |
| 1 | Participation Coefficient (Yeo7) | Age (years) | 0.9375 | 0.0500 | 0.5625 | 0.0000 |
| 2 | Participation Coefficient (Yeo7) | Age (years) | 0.8750 | 0.0500 | 0.5000 | 0.5000 |
| 3 | Participation Coefficient (Yeo7) | Age (years) | 0.6875 | 0.0000 | 0.5625 | 0.4500 |
| 4 | Participation Coefficient (Yeo7) | Age (years) | 0.0000 | 0.0000 | 0.6250 | 0.4000 |
| 5 | Participation Coefficient (Yeo7) | Age (years) | 0.6875 | 0.1500 | 0.0625 | 0.0000 |
| 1 | Participation Coefficient (Yeo7) | Motion (mm) | 0.6250 | 0.7000 | 0.6250 | 0.0000 |
| 2 | Participation Coefficient (Yeo7) | Motion (mm) | 0.5000 | 0.5000 | 0.5625 | 0.4500 |
| 3 | Participation Coefficient (Yeo7) | Motion (mm) | 0.6250 | 0.8000 | 0.0625 | 0.0000 |
| 4 | Participation Coefficient (Yeo7) | Motion (mm) | 0.2500 | 0.7000 | 0.6250 | 0.0000 |
| 5 | Participation Coefficient (Yeo7) | Motion (mm) | 0.6875 | 0.7500 | 0.8125 | 0.0500 |

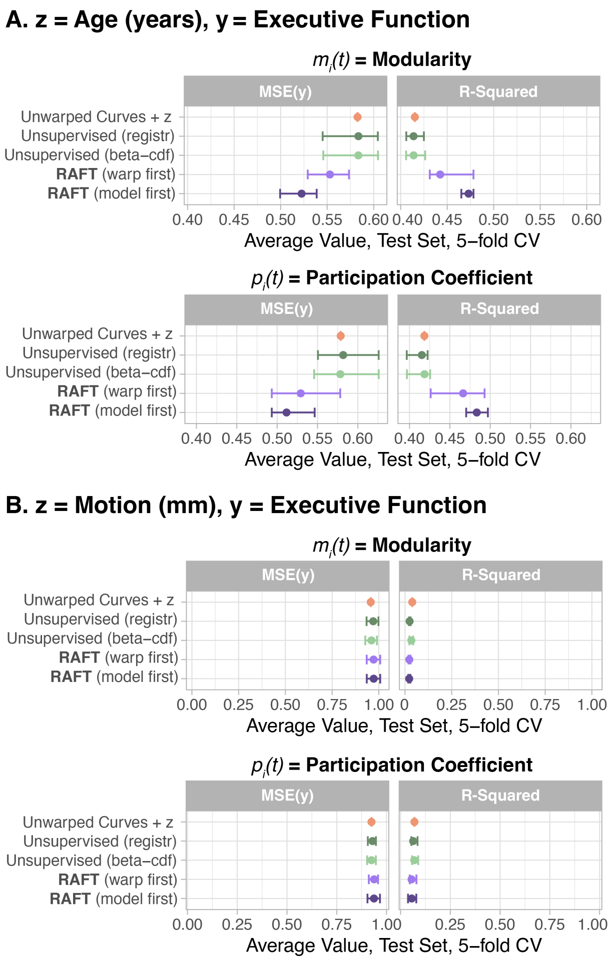


Fig. S.9. Comparison of alignment methods and unaligned curves in scalar-on-function regression models with a linear term for when $z$ reflected age (Panel A) or motion (Panel B). When $z$ reflected age, RAFT performs well against other methods. But when $z$ reflected motion, all methods perform similarly, indicating that alignment of the network diagnostic curves adds little to the predictive power beyond that given by the linear term.

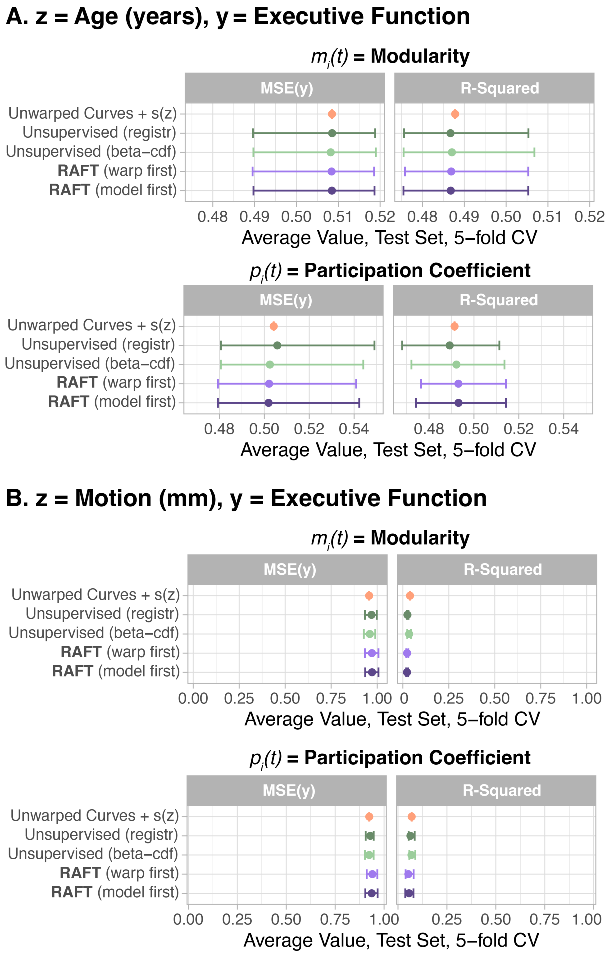


Fig. S.10. Comparison of alignment methods and unaligned curves in scalar-on-function regression models with a nonlinear term $s(z)$ for when $z$ reflected age (Panel A) or motion (Panel B). When $z$ reflected age, the average performance of all methods is similar, indicating that age-supervised alignment of the network diagnostic curves adds little to the predictive power beyond that given by the $s(z)$ term.

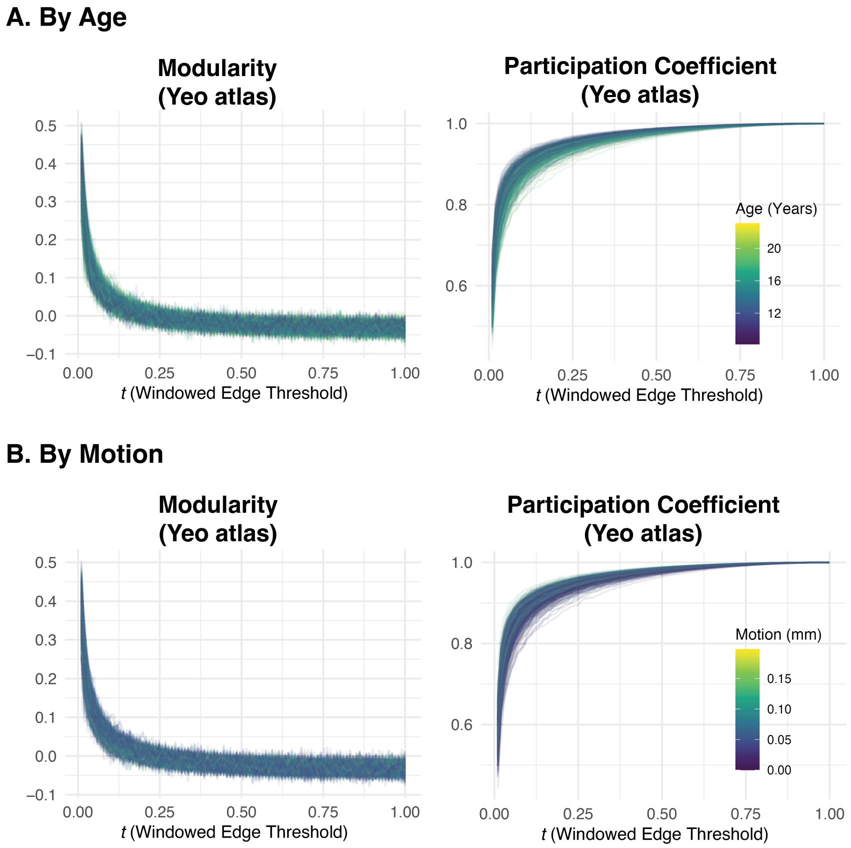


Fig. S.11. Data from the Philadelphia Neurodevelopmental Cohort (Satterthwaite *and others*, 2016) using the alternate windowed thresholding approach. Modularity and participation coefficient were calculated given the 7 communities defined in Yeo *and others* (2011) as a function of the percentile $t$, where $t = 0.01$ corresponds to the strongest percentile of edges, $t = 0.02$ corresponds to the next percentile, and so forth. Compared to those obtained from proportional thresholding, window-threshold modularity curves were noisier for $t >= 0.2$ (i.e., the weakest 80 percentile bins), likely because weaker edges have a more random configuration. In contrast, the participation coefficient curves appeared more similar for both thresholding types. The color of the curve corresponds to age (Panel A; darker = younger) or motion (Panel B; darker = less motion).

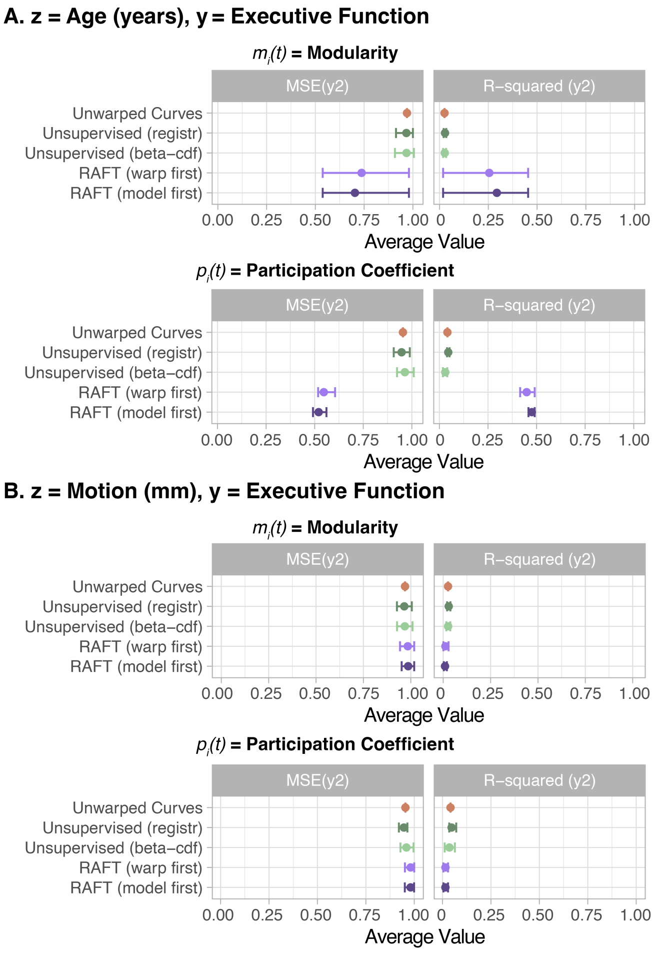


Fig. S.12. In data from the Philadelphia Neurodevelopmental Cohort (Satterthwaite *and others*, 2016), thresholded using the alternate windowed approach, RAFT alignment of the modularity and participation coefficient curves improved the prediction of executive function when the auxiliary variable was age, but not when the auxiliary variable was motion. We considered modularity (left column) and participation coefficient (right column) using communities defined by the 7-community parcellation described in (Yeo *and others*, 2011); age was measured in years, and motion was measured as average relative RMS displacement in millimeters. As with the proportional-threshold curves, RAFT performed better when $z$ reflected age (Panel A) compared to when $z$ reflected motion (Panel B), likely for similar reasons.

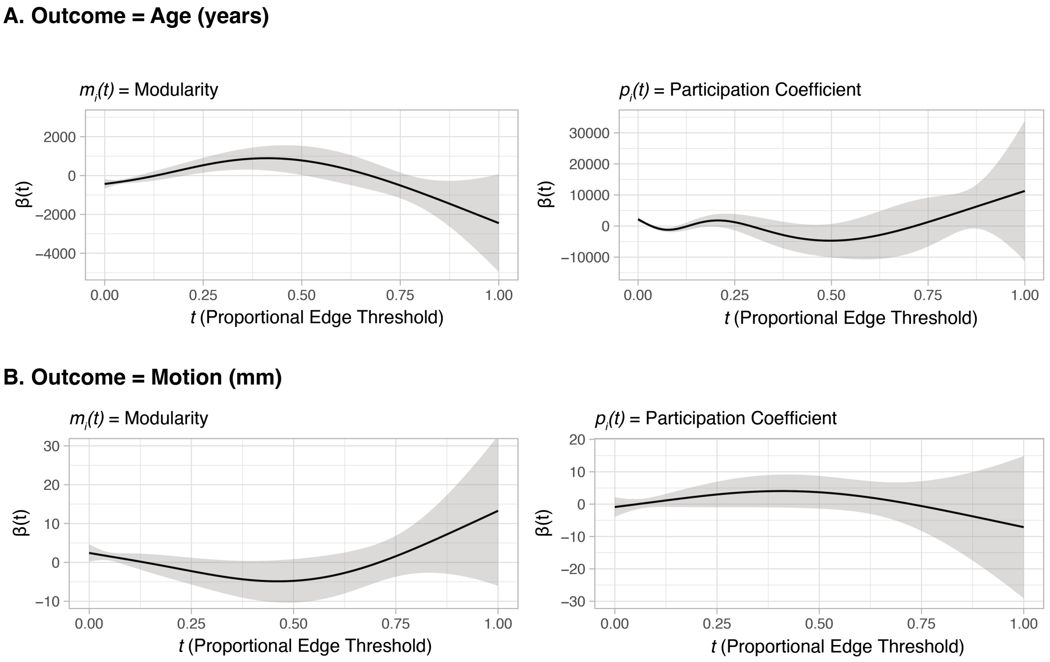


Fig. S.13. Non-linear, conditional associations between the auxiliary variables $z$ and the *unwarped* network diagnostic curves in their entirety. (Panel A) Scalar-on-function regression coefficients where age is the outcome and the predictor is either modularity (left) or participation coefficient (right). We found significant associations between age and modularity curves for $t < 0.5$, and between age and participation coefficient curves for $t < 0.3$. (Panel B) Scalar-on-function regression coefficients where motion is the outcome and the predictor is either modularity (left) or participation coefficient (right). We found significant associations between motion and modularity for $t < 0.15$, and we did not observe any significant associations between motion and participation coefficient.

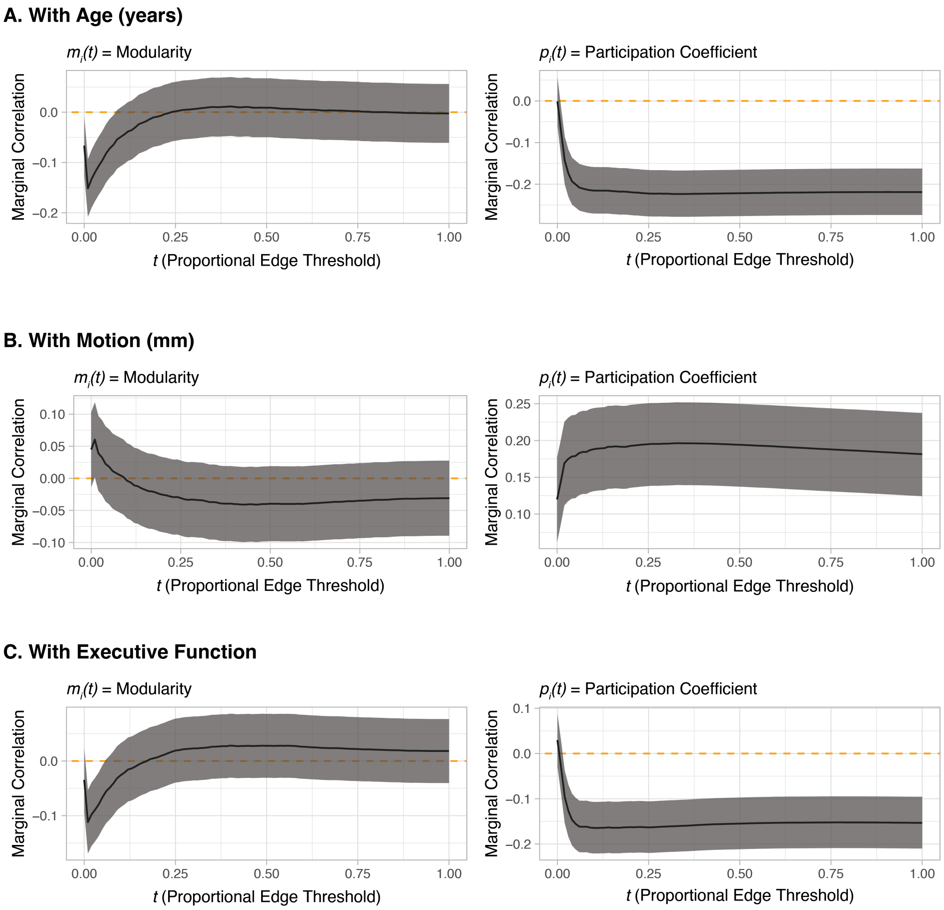


Fig. S.14. Marginal linear relationships (correlations) between the auxiliary variables $z$ or executive function $y$, and the network diagnostic curves at each threshold $t$. The correlations and their 95% percent confidence intervals were calculated separately for each value of $t$. (Panel A) The marginal correlation, as a function of $t$, between age and modularity (left), and between age and participation coefficient (right). (Panel B) The marginal correlation, as a function of $t$, between motion and modularity (left), and between motion and participation coefficient (right). (Panel C) The marginal correlation, as a function of $t$, between executive function and modularity (left), and between executive function and participation coefficient (right). This figure and Figure S.13 contain complementary information: While the *non-linear* associations in Figure S.13 are between the scalar variables and the entire curves (i.e., jointly over all $t$), these are *linear* associations between the scalar variables and the network diagnostics at a fixed value of $t$, independent of all other values of $t$.

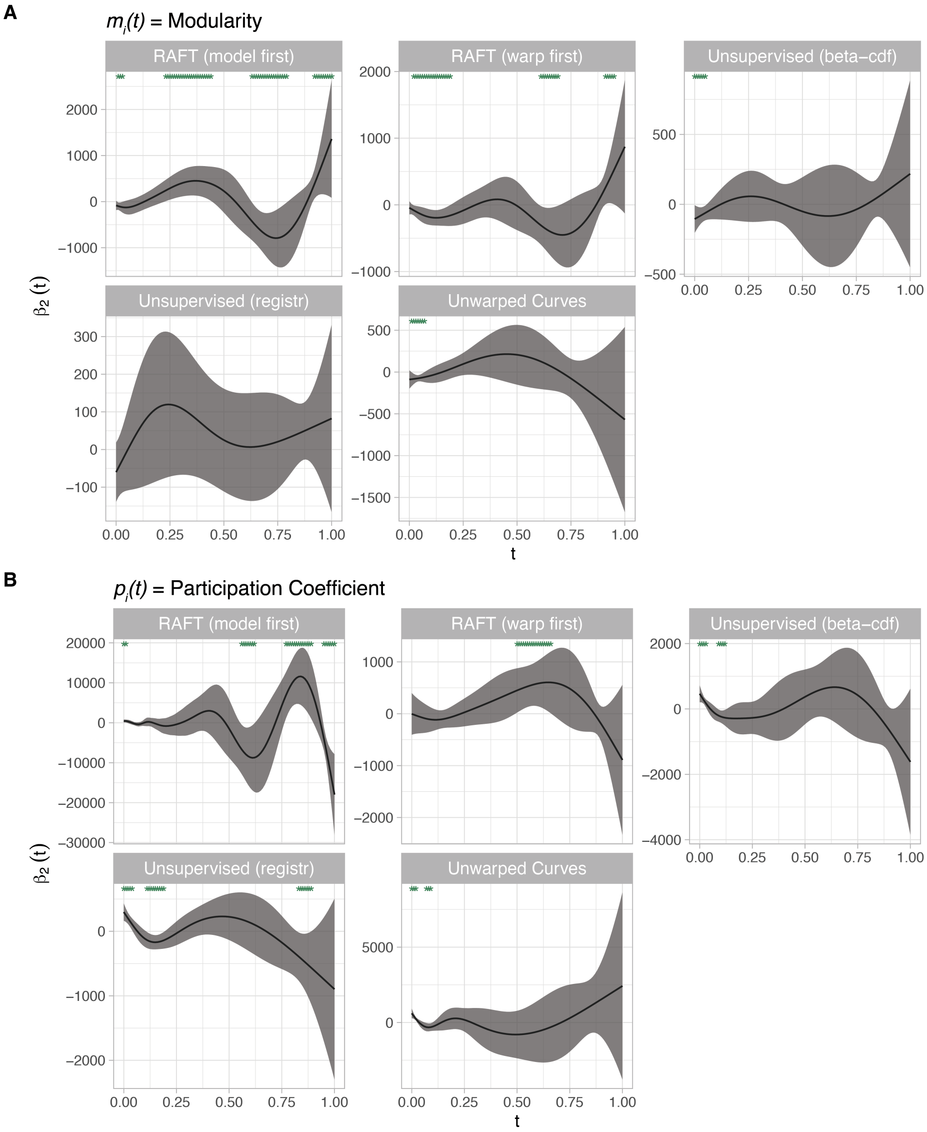


Fig. S.15. Estimated $\hat{\beta}_2(t)$ from Model 2.2 representing the association between executive function and modularity (Panel A) or participation coefficient (Panel B) after alignment by $z =$ age. The unsupervised and supervised models used the penalty parameters obtained from the cross-validation step (fold 3) provided in Table S.2. Asterisks at the top of the panel signal regions of the functional domain $t$ where $\hat{\beta}_2(t) \neq 0$. Note that these coefficient functions each correspond to a different set of aligned covariate curves.

Satterthwaite, Theodore D., Elliott, Mark A., Ruparel, Kosha, Loughead, James, Prabhakaran, Karthik, Calkins, Monica E., Hopson, Ryan, Jackson, Chad, Keefe,

Jack, Riley, Marisa, Mentch, Frank D., Sleiman, Patrick, Verma, Ragini, Davatzikos, Christos, Hakonarson, Hakon, Gur, Ruben C. *and others*. (2014, February). Neuroimaging of the philadelphia neurodevelopmental cohort. *NeuroImage* **86**, 544–553.

Schaefer, Alexander, Kong, Ru, Gordon, Evan M, Laumann, Timothy O, Zuo, Xi-Nian, Holmes, Avram J, Eickhoff, Simon B and Yeo, B T Thomas. (2017, July). Local-global parcellation of the human cerebral cortex from intrinsic functional connectivity MRI. *Cerebral Cortex* **28**(9), 3095–3114.

Tooley, Ursula A, Mackey, Allyson P, Ciric, Rastko, Ruparel, Kosha, Moore, Tyler M, Gur, Ruben C, Gur, Raquel E, Satterthwaite, Theodore D and Bassett, Danielle S. (2019, April). Associations between neighborhood SES and functional brain network development. *Cerebral Cortex* **30**(1), 1–19.

Yeo, B. T. Thomas, Krienen, Fenna M., Sepulcre, Jorge, Sabuncu, Mert R., Lashkari, Danial, Hollinshead, Marisa, Roffman, Joshua L., Smoller, Jordan W., Zöllei, Lilla, Polimeni, Jonathan R., Fischl, Bruce, Liu, Hesheng *and others*. (2011, September). The organization of the human cerebral cortex estimated by intrinsic functional connectivity. *Journal of Neurophysiology* **106**(3), 1125–1165.
